# A bifunctional imprinted lncRNA scaffolds NuRD–HDAC1 to couple cis-activation of Dio3 with trans-repression of osteoblast chromatin

**DOI:** 10.1101/2025.04.14.647401

**Authors:** Yuechuan Chen, Benjamin Wildman, Mohammad Rehan, Snehasis Mishra, Shazia Khan, Lubana H. Afreen, Quamarul Hassan

## Abstract

Long non-coding RNAs (lncRNAs) frequently act at imprinted loci, yet how a single lncRNA can simultaneously activate its neighbor and repress distant genes through a shared chromatin platform remains poorly defined. We show that imprinted Dio3os performs this bifunctional role by scaffolding the nucleosome remodeling and histone deacetylase (NuRD–HDAC1) complex. ATAC-seq and ChIP-seq across osteoblast differentiation reveal that the Dio3–Dio3os bidirectional promoter is dynamically remodeled and co-occupied by HDAC1, HIF1α, and active histone marks. CRISPR-mediated exon deletion, CRISPRi, and polyadenylation termination demonstrate that Dio3os activates Dio3 in cis while repressing osteogenic genes in trans. RIP–MS and reverse pulldown identify direct Dio3os–NuRD interaction; ChIP shows the same machinery produces opposing H3K27ac outcomes at Dio3os versus osteogenic promoters. In vivo, osteoblast-specific CRISPRi increases trabecular and cortical bone mass, whereas CRISPRa reduces it in both sexes. Dio3os couples chromatin remodeling to thyroid-hormone metabolism through a context-dependent NuRD scaffold.

**One sentence summary:** *Dio3os* scaffolds NuRD–HDAC1 to activate *Dio3* in cis and repress thyroid-hormone–responsive osteogenic genes in trans.

## INTRODUCTION

Long noncoding RNAs (lncRNAs) act as flexible scaffolds that bring chromatin-remodeling and transcription complexes to specific genomic addresses, enabling tissue- and stage-specific gene regulation (^1–5^). At imprinted loci in particular, lncRNAs are central to coordinated regulation of co-expressed gene clusters (^6–12^). A long-standing question is whether and how a single lncRNA can produce two opposite transcriptional outputs, local activation and distant repression, through a shared protein platform. Resolving this question is critical for understanding how the same chromatin machinery (e.g., NuRD, PRC2, HDAC complexes) can support context-specific gene regulation, with implications for development, hormone signaling, and disease.

Thyroid hormones (T3, T4) are essential regulators of skeletal development and homeostasis (^13, 14^); imbalances cause well-recognized bone disease, and the prevalence of hypothyroidism in adults continues to rise (^14–17^). Intracellular thyroid-hormone availability is set by three deiodinases: DIO1 and DIO2 activate T4 to T3, whereas DIO3 inactivates T3/T4 to rT3/T2 (^18, 19^). DIO3 is broadly expressed in skeletal lineages and is essential for bone health, as shown by perinatal thyrotoxicosis, adult hypothyroidism, and growth defects in *Dio3*-null mice (^20–23^). Consumptive hypothyroidism, a rare pediatric condition driven by aberrant *DIO3* expression, illustrates the clinical consequences of dysregulated DIO3 (^24, 25^). Despite this, we still do not understand how DIO3 expression is coordinated with osteoblast-specific thyroid-hormone signaling. The *DLK1–DIO3* locus is one of the best-characterized imprinted regions in mammalian genomes and contains numerous noncoding RNAs (^26–28^). *Dio3os,* transcribed antisense to Dio3 from a shared bidirectional promoter, contributes to multiple cancers (^29–31^), and GWAS meta-analyses have identified it as a thyroid-hormone–associated locus, but its mechanism of action remains unknown. Most published lncRNA mechanisms describe a single output either local cis regulation (e.g., *Airn*, *Kcnq1ot1*) or trans-acting scaffolding (e.g., *HOTAIR*, *ANRIL*) (^6, 7, 32, 33^). Whether a single mammalian lncRNA can integrate both modes through a defined chromatin platform, and what determines its local-versus-distal specificity, remain open questions.

Here, using multiple CRISPR-based perturbations that preserve *Dio3* coding integrity, we show that *Dio3os* acts as a bifunctional regulator: it activates *Dio3* in cis through promoter recruitment of HDAC1/HIF1α-dependent NuRD, while repressing osteogenic gene programs in trans through the same NuRD–HDAC1 complex. We map this activity to specific *Dio3os* exons, identify the NuRD complex as the molecular interactor, demonstrate context-dependent H3K27ac output at *Dio3os* versus osteogenic loci, and establish bidirectional in vivo causality with osteoblast-specific CRISPRi and CRISPRa in both sexes. The findings define an imprinted lncRNA–NuRD–HDAC1 axis that couples chromatin remodeling to thyroid-hormone metabolism, with implications across lncRNA biology, endocrine regulation, and skeletal disease.

## RESULTS

### Mouse and human Dio3os share conserved bidirectional promoter architecture

To identify lncRNAs that change during osteogenic differentiation, we profiled 80 evolutionarily conserved lncRNAs in mouse MC3T3-E1 osteoblasts at days 3, 6, 9, and 11 using a human- specific lncRNA qPCR library probed with murine cDNAs (Fig. 1A, 1B). Among differentially expressed lncRNAs, *Dio3os*, *lincRNA-p21*, *antisense Nespas*, and *Emx2os* all transcriptionally share a bidirectional promoter architecture with an upstream coding partner (*Dio3*, *p21*, *Nesp*, *Emx2*; Fig. 1C), allowing transcriptional changes in cis to be distinguished from trans effects. We focused on the *Dio3os* locus because of its candidate role in hypothyroidism-associated skeletal disease. Consistent with prior reports, the *Dio3–Dio3os* intergenic region functions as a bidirectional regulatory element rather than a single defined 1.2-kb promoter (17). The transcripts overlap at their 5’ ends within a G+C–rich region, a feature characteristic of imprinted antisense-promoter activity. In the mouse, *Dio3os* transcript variant 203 (1387 bp, four exons) spans ∼3 kb and shares a 1.2-kb bidirectional promoter with *Dio3* (Fig. 1D). In humans, *DIO3OS* variant 203 (1156 bp, four exons; Fig. 1E) and variant 205 (intron less, 3128 bp; Fig. 1F) preserve this organization. An 82-nt (mouse) or 136-nt (human) miR-1247 is embedded in exon 1 of both. Knockdown of miR-1247-3p and -5p in MC3T3-E1 cells (∼70% depletion) had minimal effect on *Dio3os*, *Dio3*, or osteogenic markers (fig. S1A, S1B), indicating that the embedded microRNA is not a major driver of the *Dio3os* function characterized in this study.

**Figure 1:**
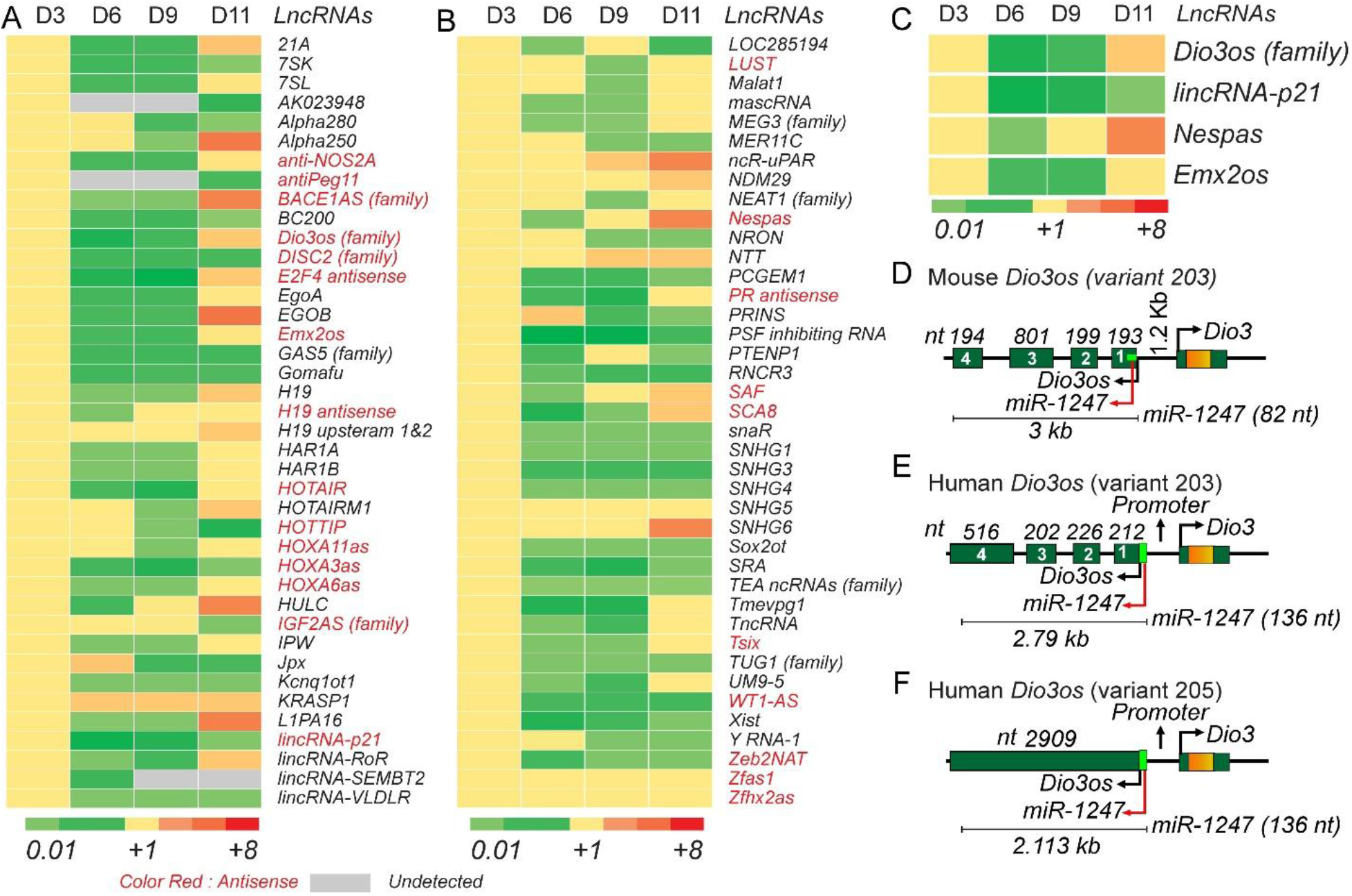
Differential expression of lncRNAs during murine osteogenic differentiation using human LncProfiler qPCR arrays. (A and B) Heatmap showing 80 differentially expressed mouse lncRNAs during osteogenic differentiation of MC3T3-E1 cells at Days 3, 6, 9, and 11 when probed with human-specific lncRNA qPCR arrays. The total RNA of MC3T3-E1 cells during differentiation time points was used for LncRNA analysis. Relative fold changes of the LncRNAs were hierarchically clustered by using dChip software. Expression values are represented as log2 fold changes, with upregulated lncRNAs indicated in red and downregulated lncRNAs in green. The red color highlights antisense transcripts. We performed three independent LncProfiler replicates at osteoblast differentiation time points (n = 3) to ensure the reproducibility of our LncProfiler findings. (C) A heatmap showed four significantly changed sense and antisense or opposite-strand LncRNAs, including Dio3os, which shared a bidirectional promoter during osteogenesis. Comparison of the Structure of Dio3os between Humans and Mice. (D) Schematic of the Dio3os-Dio3 locus structure between mouse and human reveals that the mouse Dio3os transcript (variant 203) contains four exons (E1–E4) with no ATG codon in the first exon, spanning approximately 3 kb in total. (E&F) In humans, there are two Dio3os transcript variants (203 and 205). The 2.79 kb human variant 203 transcript also contains four exons, similar to the mouse gene. However, the human variant 205 consists of a single long exon spanning approximately 3.1 kb. The intergenic miR-1247 is highlighted with a red arrow, with its size indicated, and the 5’ and 3’ UTR regions are also marked. Exon sizes (in nucleotides) are labeled for comparison across species.

### Chromatin accessibility, histone marks, and osteogenic transcription factor RUNX2 occupancy define stage-specific promoter regulation

To elucidate the chromatin state of the murine bidirectional promoter at chromosome 12, we analyzed chromatin accessibility (ATAC-seq) in MC3T3-E1 osteoblasts and histone modifications (ChIP-seq) in bone marrow stromal cells (BMSCs) during the osteogenic differentiation process (Fig. 2A). ATAC-seq revealed high accessibility at the 1.2-kb *Dio3os* promoter and exons 2–4 during proliferation, declining at early differentiation (day 7; Fig. 2B). ChIP-seq across days 0, 7, 14, and 21 showed H3K4me3 and H3K27ac enrichment at the promoter during proliferation that declined with differentiation (Fig. 2C, 2D). The H3K36me3 modifications in the gene body were correspondingly elevated during proliferation, consistent with active transcription (Fig. 2F). The repressive H3K27me3 mark was low at the promoter but enriched upstream of *Dio3* (Fig. 2E), suggesting fine-tuning of *Dio3* output. By days 14 and 21, both active and repressive marks diminished, indicating a poised rather than permanently silenced state (^34–36^). RUNX2 binding at the *Dio3–Dio3os* promoter increased markedly from days 7 through 21 (Fig. 2G), coinciding with the loss of H3K4me3/H3K27ac and consistent with the RUNX2-mediated repression of promoter activity that we observe in reporter assays (Fig. 3D, 3E). Across 18 kb of the *Dio3os* locus, H3K4me1 and H3K27ac enrichment were minimal in the intragenic region (exons 2–4). In contrast, the promoter-proximal region showed strong H3K4me3 and H3K27ac (fig. S2), indicating that *Dio3os* regulation is promoter-driven rather than enhancer-dependent (^37, 38^).

**Figure 2:**
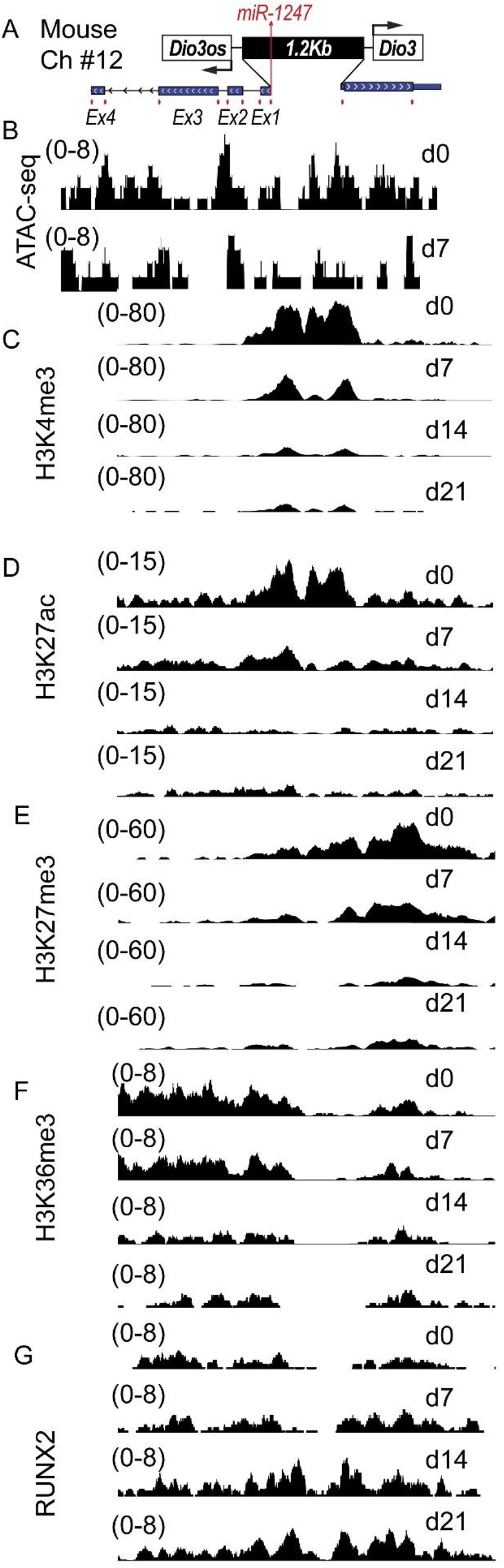
Chromatin accessibility and histone modifications at the 1.2 kb mouse Dio3os-Dio3 promoter during osteoblast differentiation. (A) Schematic of the Dio3-Dio3os promoter, miRNA 1247, and exonic regions at chromosome 12. (B) Chromatin accessibility profiles at the Dio3-Dio3os gene loci. Accessibility was assayed on Day 0 during MCM3T3-E1 proliferation and on Day 7 at the onset of differentiation. The “Y-axis” indicates accessible genomic tracts, and the “X-axis” denotes the genomic regions. (C, D, E, F, & G) BMSCs isolated from mouse bone marrow, differentiated under osteogenic conditions, and ChIP-sequencing was performed using H3K4me3, H3K27ac, H3K27me3, H3K36me3, and RUNX2 antibodies at days 0, 7, 14, and 21 to understand the chromatin modification status of the 1.2 kb Dio3-Diosos bidirectional promoter during BMSC differentiation. “Y-axis” indicates genomic tracts showing enrichment for H3K4me3, H3K27ac, H3K27me3, H3K36me3, and RUNX2 at promoters, and the “X” axis denotes the genomic regions. In the promoter, quantitative H3 modification peaks of K4me3 and K27ac indicate active regions, K27me3 indicates repressive regions, while K36me3 modification peaks in the gene body indicate gene synthesis levels. At each time point, we prepared two independent biological replicates of ChIP-Seq libraries, along with two input libraries derived from sonicated DNA. For ATAC-seq, at each time point (days 0 and 7), we prepared independent ATAC-seq libraries and their corresponding input libraries, generated from DNA treated with Tn5 transposase.

**Figure 3:**
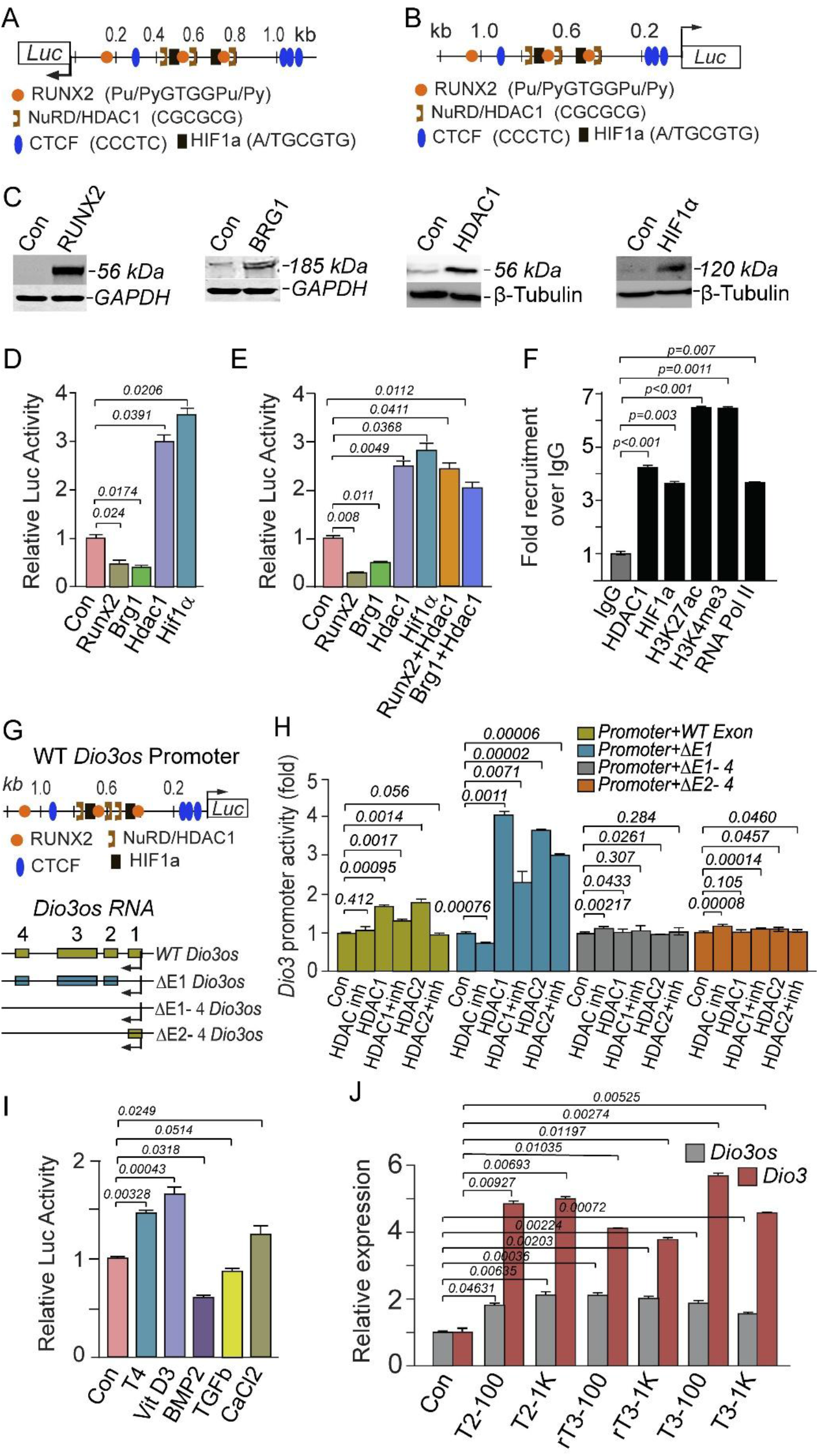
Dio3–osteogenic factors coordinately regulate Dio3os promoter activity and Dio3os exon–dependent HDAC1/2 function. (A) The luciferase reporter is positioned downstream of the Dio3os transcriptional direction. (B) The reporter is placed downstream of the Dio3 transcriptional direction to measure bidirectional promoter activity. (C) Western blot analysis shows increased levels of RUNX2, BRG1, HDAC1, and HIF1α. (D) Relative luciferase activity of the Dio3os promoter was measured in cells overexpressing Runx2, Brg1, Hdac1, and Hif1a. (E) Relative luciferase activity of the Dio3 promoter was assayed with RUNX2, Brg1, Hdac1, and Hif1a alone, or with RUNX2+Hdac1 and Brg1+Hdac1 together. All respective overexpression plasmids were in mammalian expression vectors. (F) Chromatin immunoprecipitation (ChIP) was performed using antibodies against HDAC1, HIF1α, H3K27ac, H3K4me3, and RNA polymerase II. Enrichment at the Dio3–Dio3os promoter is shown as fold recruitment over IgG control. Data are presented as mean ± SEM (n = 3). Statistical significance was determined relative to the IgG control as indicated. (G, upper panel) Bioinformatics analysis of the promoter nucleotide sequences identified several key regulatory elements within the Dio3os-Dio3 promoter, including RUNX2, CTCF, NuRD, and HIF1a. (G, lower panel) A schematic representation of the Dio3os exon knockout models: WT (wild-type), Δ1 (Exon 1 deletion), Δ1-Δ4 (Exon 1-4 deletion), and Δ2-Δ4 (Exon 2-4 deletion). (H) Dio3-Dio3os WT promoter was transfected with four different Dio3os constructs (WT, Δ1, Δ1-Δ4, and Δ2-Δ4) and cotransfected with Hdac1 and 2 overexpression clones and treated with HDAC1 and 2 inhibitors. Control 1 (Dio3-Dio3os Promoter + WT+ Empty Vector), Control 2 (Dio3-Dio3os Promoter + WT Dio3os + HDAC1+2 inhibitor). Experimental: HDAC1 (Dio3-Dio3os Promoter + Hdac1 overexpression), HDAC1 + inh (Dio3-Dio3os Promoter + Hdac1 overexpression with inhibitor), HDAC2 (Dio3-Dio3os Promoter + Hdac2 overexpression), HDAC2 + inh (Dio3-Dio3os Promoter + Hdac2 overexpression with inhibitor), +/- WT, Δ1, Δ1- Δ4, Δ2- Δ4 constructs. We performed three independent experimental replicates (n=3) to ensure the reproducibility of our findings of HDAC1 and 2 response. P values are indicated as: NS (P > 0.05), * (P ≤ 0.05), ** (P ≤ 0.01), *** (P ≤ 0.001), and **** (P ≤ 0.0001). (I)The Dio3 promoter activities were determined with various factors, including thyroid hormone (T4, 100 ng/ml), BMP2 (100 ng/ml), and TGFβ (100 ng/ml) for 48 hours. The relative luciferase activities were normalized with promoter-less mammalian vector encoding wild-type Renilla luciferase (pRL-null). (J) qRT-PCR validation of Dio3os and Dio3 in response to increasing T2, reverse T3 (rT3), and T3 with 100 ng and 1000 ng/ml concentrations. The following iodothyronines were used in our experiment: T4 (3,3′,5,5′-tetraiodo-L-thyronine; Cat. T1775), T3 (3,3′,5-triiodo-L-thyronine; Cat. T2877), rT3 (3,3′,5′-triiodo-L-thyronine; Cat. T075), and T2 (3,3′-diiodo-L-thyronine; Cat. 719536). Relative expressions are normalized to the housekeeping gene GAPDH and represented as mean ± SEM (n = 3 replicates). We accomplished three independent experimental replicates of Luciferase reporter assays (n = 3) to ensure the reproducibility of our bidirectional promoter activity. P values are indicated as: NS (P > 0.05), * (P ≤ 0.05), ** (P ≤ 0.01), *** (P ≤ 0.001), and **** (P ≤ 0.0001).

### HDAC1 and HIF1α activate the bidirectional promoter, and activation requires Dio3os exons 2–4

The 1.2-kb GC-rich bidirectional promoter contains a TSS in *Dio3os* exon 1 (CGCGGGAGCGCT) and a shared TATA box (TATAAAT), with four CTCF, three NuRD/HDAC1 (CGCGCG), three RUNX2 (Pu/PyGTGGPu/Py), and two HIF1α (A/TGCGTG) binding sites (Fig. 3A, 3B). We cloned the 1.2-kb promoter in both orientations into luciferase reporters to assess bidirectional activity. Western blots confirmed overexpression of RUNX2, BRG1, HDAC1, and HIF1α (Fig. 3C; fig. S7). RUNX2 and BRG1 reduced both forward and reverse promoter activity by approximately 50%. In contrast, HDAC1 and HIF1α significantly increased promoter activity by 2.5 to 3.5 times (Fig. 3D and 3E). Combined RUNX2/BRG1/HDAC1 only modestly reduced HDAC1-driven activation, indicating that HDAC1 dominance is not relieved by these repressors (Fig. 3E). ChIP-qPCR confirmed enrichment of HDAC1, HIF1α, H3K27ac, H3K4me3, and RNA polymerase II at the endogenous *Dio3–Dio3os* promoter (Fig. 3F), consistent with an active state co-occupied by HDAC1 and HIF1α.

To localize the HDAC1-interacting element within *Dio3os*, we engineered CRISPR/Cas9-mediated exon-deletion constructs (ΔE1, ΔE1–4, ΔE2–4; Fig. 3G) and tested their effect on bidirectional promoter activity in the presence of HDAC1/2 overexpression. WT *Dio3os* supported ∼1.8–2-fold HDAC-dependent activation; ΔE1 supported ∼3–4-fold activation, indicating that exon 1 is dispensable for HDAC1/2-driven promoter activation. By contrast, ΔE1–4 or ΔE2–4 abolished HDAC1/2 responsiveness (Fig. 3H). This identifies that Dio3os exons 2–4 as the functional element required for NuRD/HDAC1/2-mediated activation of the bidirectional promoter. Thyroid hormone (T4), vitamin D3, and CaCl2 increased promoter activity; BMP2 and TGFβ decreased it (Fig. 3I). Direct treatment with T2, rT3, or T3 (100 or 1000 ng/ml) modestly induced *Dio3os* and strongly induced *Dio3* (>3-fold; Fig. 3J), confirming that iodothyronines drive transcription in both directions of the bidirectional promoter.

### Dio3os activates Dio3 in cis and represses osteogenic genes in trans

To resolve cis from trans effects, we combined gain-of-function (CMV-driven overexpression) with multiple loss-of-function approaches (CRISPR/Cas9 exon deletion, dCas9–KRAB CRISPRi, and engineered polyadenylation termination) in pre-osteoblast MC3T3-E1 cells. Overexpression of *Dio3os* for 24 hours increased *Dio3* mRNA, while *Runx2* and *Ocn* were reduced by 70–80% (Fig. 4A), consistent with cis-activation of *Dio3* and trans-repression of osteogenic genes. CRISPR/Cas9 deletion of exon 1 (ΔE1) or exons 1–4 (ΔE1–4) was verified by PCR genotyping, Sanger sequencing, and genomic BLAST (fig. S3). ΔE1–4 reduced *Dio3os* by >90% and *Dio3* by ∼80% without affecting the *Dio3* coding sequence, and induced *Runx2* 4-fold and *Ocn* 8-fold (Fig. 4B). ΔE1 alone reduced *Dio3* and *Dio3os* but did not alter *Runx2* or *Ocn*, indicating that the trans-repressive activity requires sequences in exons 2–4. ΔE1–4 cells showed strong ALP staining at day 14 and enhanced mineralization (Von Kossa) at day 21 relative to control (Fig. 4C; additional replicates in figs. S4C and S4D). CRISPRi targeting *Dio3os* between exons 1 and 2 reduced *Dio3os* and *Dio3* while upregulating *Runx2* and *Ocn* (Fig. 4D), confirming both cis and trans functions in a transcript-preserving setting. Insertion of polyadenylation (pAS) sequences between exons 1–2, 2–3, or 3–4 terminated *Dio3os* transcription at distinct positions while preserving promoter integrity. Each pAS insertion reduced *Dio3os* and *Dio3* and upregulated *Ocn* and *Runx2* 4–20-fold (Fig. 4E). Four independent perturbation strategies, including overexpression, exon deletion, CRISPRi, and transcription termination, thus converge on the same dual function: cis-activation of *Dio3* and trans-repression of osteogenic genes.

**Figure 4:**
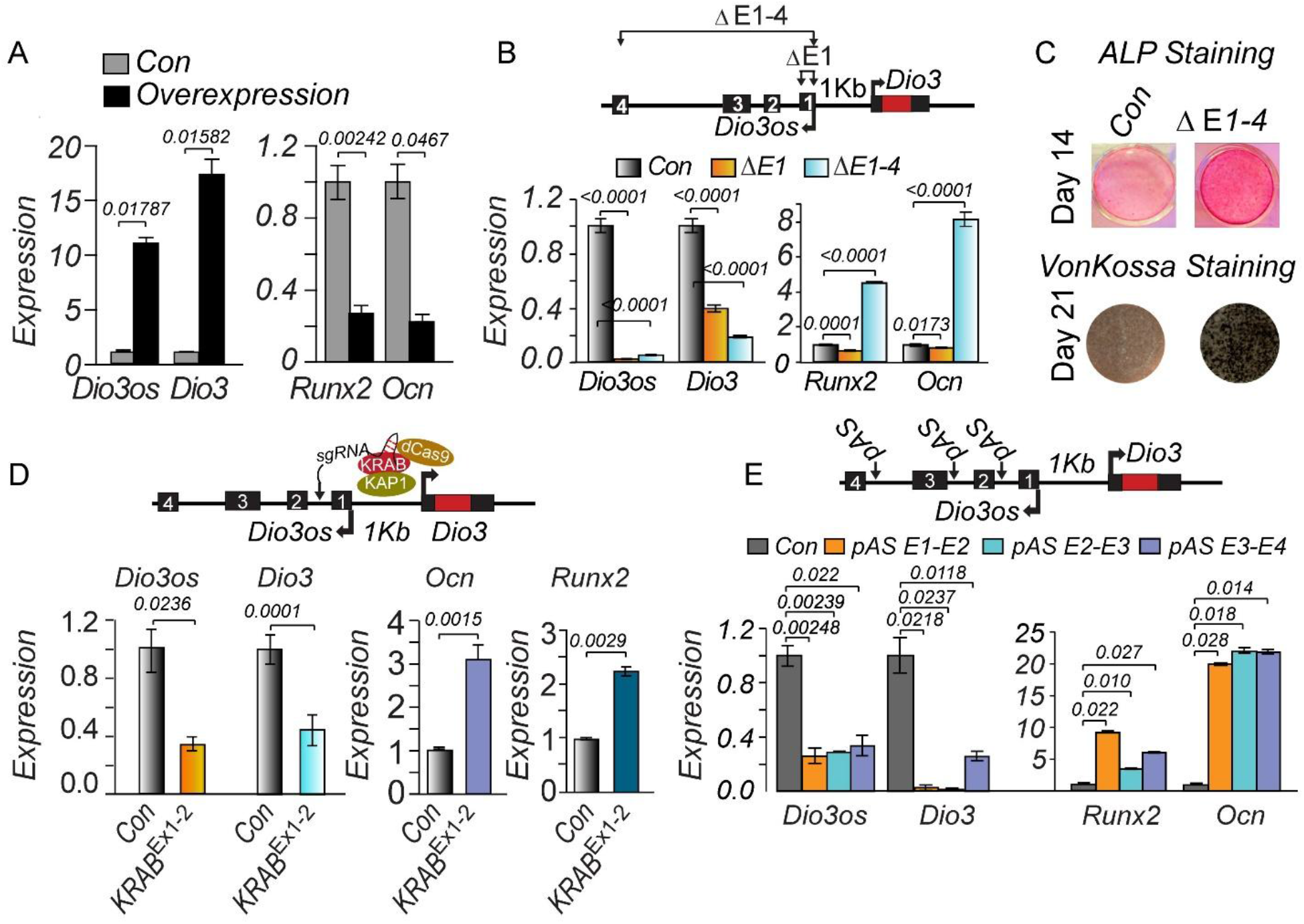
Dio3os gain of function, CRISPR-mediated Dio3os knockout, CRISPR inhibition, and exon polyadenylation indicate that Dio3os function is crucial for osteoblast differentiation and maturation. (A) Quantitative mRNA and RNA analysis of Dio3os, Dio3, Runx2, and Ocn upon overexpression of Dio3os. (B) CRISPR-Cas9-mediated knockout of Dio3os exons using exon-specific sgRNA. ΔEx1: Exon 1 (90 nt); 2) ΔEx1-4: Exons 1-4, including the intronic regions (2.7 kb). Lower panel: lncRNA and mRNA levels for Dio3os, Dio3 (left), and Runx2 and Ocn (right). (C) Images of alkaline phosphatase (ALP) staining and Von Kossa staining (lower panel) of CRISPR-Cas9 exons 1-4 knocked-out cells with Dio3os-specific sgRNA compared to non-specific sgRNA, confirming osteoblast activity and matrix mineralization. (D) CRISPRi targeting of the Dio3os exon 1–2 region using dCas9-KRAB/KAP1 represses Dio3os and Dio3 expression and increases osteogenic markers Ocn and Runx2. Data are mean ± SEM (n = 3); p values by two-tailed Student’s t-test. (E) Upper panel: Insertion of polyadenylation signal at the end of the first (-621 nt), second (-1546 nt), and third exons (-2784 nt) of Dio3os gene using CRISPR-Cas9. Lower panel: lncRNA and mRNA levels for Dio3os, Dio3 (left), and Runx2 and Ocn (right). We performed three independent experimental replicates (n = 3) of Dio3os overexpression, exons CRISPR knockout, ALP and Von Kossa staining, and exons polyadenylation to confirm and validate the reproducibility of Dio3os expression and function. P values are indicated as: NS (P > 0.05), * (P ≤ 0.05), ** (P ≤ 0.01), *** (P ≤ 0.001), and **** (P ≤ 0.0001) with 3 replicates.

### Dio3os loss derepresses thyroid-hormone–responsive osteogenic genes and opens their promoters

To uncover the role of Dio3os in the process of initiation and maturation of osteoblast differentiation, we performed integrated RNA-seq (day 0) and ATAC-seq (day 14) in WT and Dio3os knockout (ΔE1–4) MC3T3-E1 osteoblasts. Sequencing libraries had >97% total and >84% uniquely mapped reads (fig. S4A), with high replicate concordance (R² = 0.957; fig. S4B). ATAC-seq confirmed loss of *Dio3os* exonic accessibility (exons 2-4), with reduced transcription of *Dio3os* and *Dio3* (Fig. 5A, 5B). RNA-seq revealed 285 upregulated and 367 downregulated genes in ΔE1–4 (Fig. 5B). GO enrichment of upregulated transcripts was dominated by ossification, bone mineralization, and osteoblast differentiation (Fig. 5C). Twenty-two thyroid-hormone–responsive osteogenic genes, including *Runx2*, *Sp7*, *Col1a1*, *Alpl*, and *Ocn,* increased ≥2-fold (Fig. 5D). The non-TH-responsive Mmp9 was reduced, possibly through non-genomic thyroid-hormone signaling via integrin αvβ3 (^39, 40^). Rpl13a and Cdk6 acted as controls that were not responsive to thyroid hormones (Fig. 5D, 5E). ATAC-seq showed accessibility gains at the promoters of the 22 TH-responsive genes (Fig. 5E), establishing chromatin opening as the basis for the transcriptional changes. *Dio2* promoter accessibility is low in WT during early differentiation (^14, 41^). It is increased in ΔE1–4 cells without T4 stimulation (Fig. 5E), consistent with a feedback mechanism in which reduced *Dio3* lowers thyroid hormone inactivation, prompting *Dio2* upregulation to restore T3 balance (^22,23,42^).

**Figure 5:**
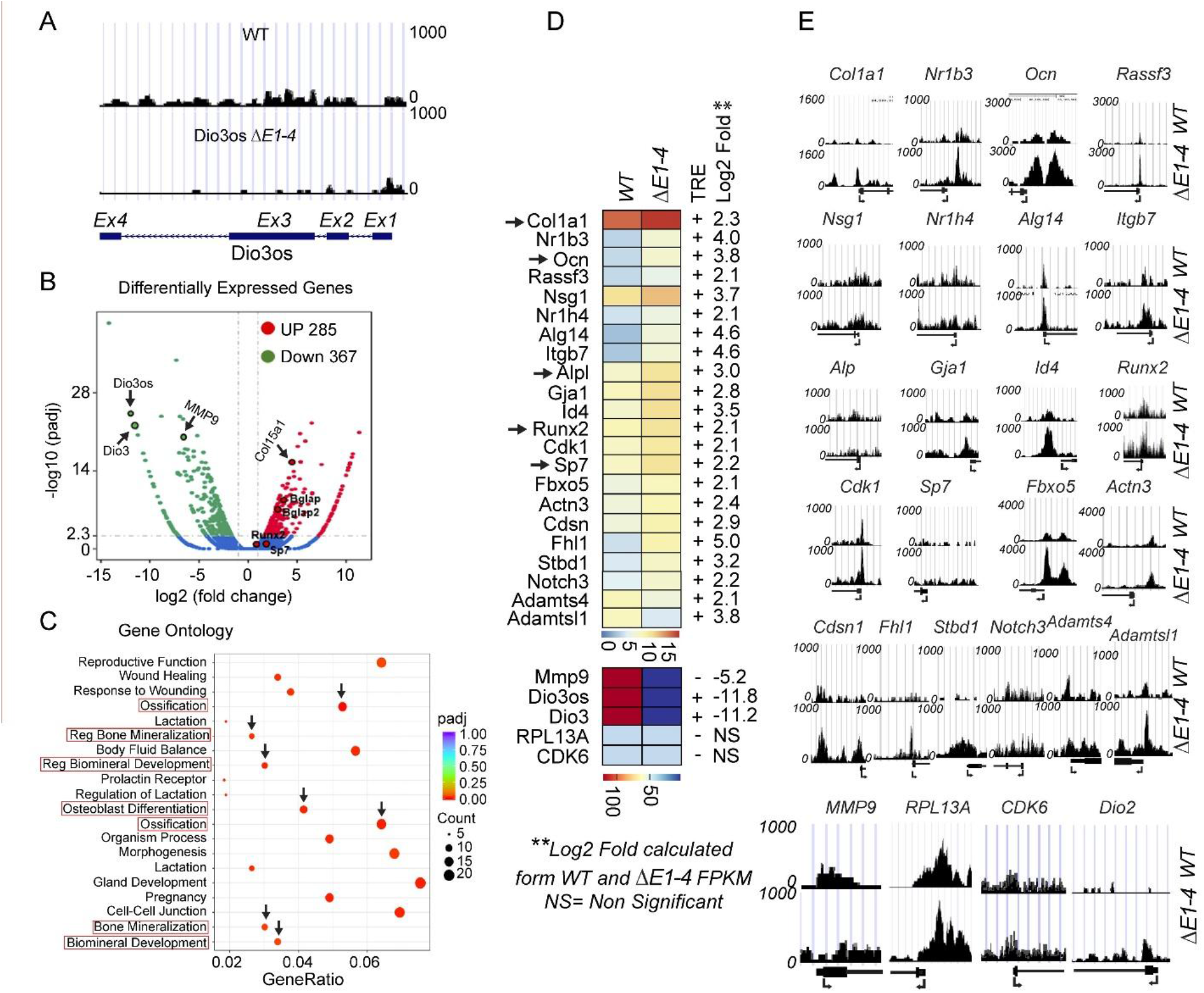
Integrated RNA-seq and chromatin accessibility analyses reveal Dio3os-dependent transcriptional regulation of thyroid hormone–responsive osteogenic genes. (A) Genome browser tracks showing ATAC-seq reads across the Dio3os–Dio3 locus in wild-type (WT) and Dio3os knockout (ΔE1–4) osteoblasts, indicating loss of Dio3os exonic openness (B) Volcano plot of differentially expressed genes (DEGs) in Dio3os KO versus WT cells showing 285 upregulated and 367 downregulated transcripts (log₂ fold change ≥ 1, padj < 0.05); representative Dio3os- regulated genes are labeled. (C) Gene Ontology (GO) enrichment analysis of upregulated genes highlights processes related to ossification, bone mineralization, and osteoblast differentiation. (D) Heatmap showing the top 22 thyroid hormone–responsive genes upregulated in Dio3os KO cells, including Col1a1, Ocn, Runx2, and Sp7; fold changes are expressed as log₂(FPKM KO/WT). Thyroid hormone cis-regulatory elements were identified using Ensembl and TRANSFAC bioinformatics tools. The color scale is based on the relative fold change in expression. (E) Genomic tracks for ATAC-seq data were generated using the standard protocol. X-axis, genomic coordinates; Y-axis, normalized ATAC-seq read counts. Gene names are shown at the top, and transcription direction is shown at the bottom. The RNA-seq and ATAC-seq data were aligned to the mm10 mouse genome. One RNA-seq analysis with 41-53 million uniquely mapped reads was analyzed for both wild-type and Dio3os knockout samples.

Genome-wide ATAC analysis confirmed a global increase in chromatin accessibility around transcription start sites in ΔE1–4 cells, concentrated within ≤1 kb of promoters with minimal change in distal regions (fig. S5A, S5B). ALP staining at day 14 confirmed enhanced osteogenic potential in ΔE1–4 cells (fig. S5C). *Dio3os* thus acts as a trans-acting chromatin regulator that coordinates thyroid-hormone metabolism with osteogenic gene expression.

### Dio3os scaffolds the NuRD–HDAC1 complex and produces context-dependent H3K27ac modifications output

To define the protein interactome that mediates *Dio3os* function, we performed RNA immunoprecipitation coupled to mass spectrometry (RIP–MS) with biotinylated *Dio3os* probes in MC3T3-E1 cells. Hits included the NuRD components HDAC1, CHD3, MBD3, RBBP7, MTA2, and the GATAD2 paralogs P66A/P66B, as well as the REST co-repressor SIN3B (Fig. 6A). Peptide-count enrichment was specific: CHD3 (2.6-fold), CHD4 (4.36-fold), MTA1 (3.79-fold), MBD3 (1.4-fold), and HDAC1 (7-fold) over IgG controls (Fig. 6B). Reverse pulldown using antibodies against HDAC1, MBD3, and MTA1 confirmed robust endogenous *Dio3os* enrichment (22-, 15-, and 7.5-fold over IgG, respectively; Fig. 6C), establishing direct association between *Dio3os* and core NuRD components.

**Figure 6:**
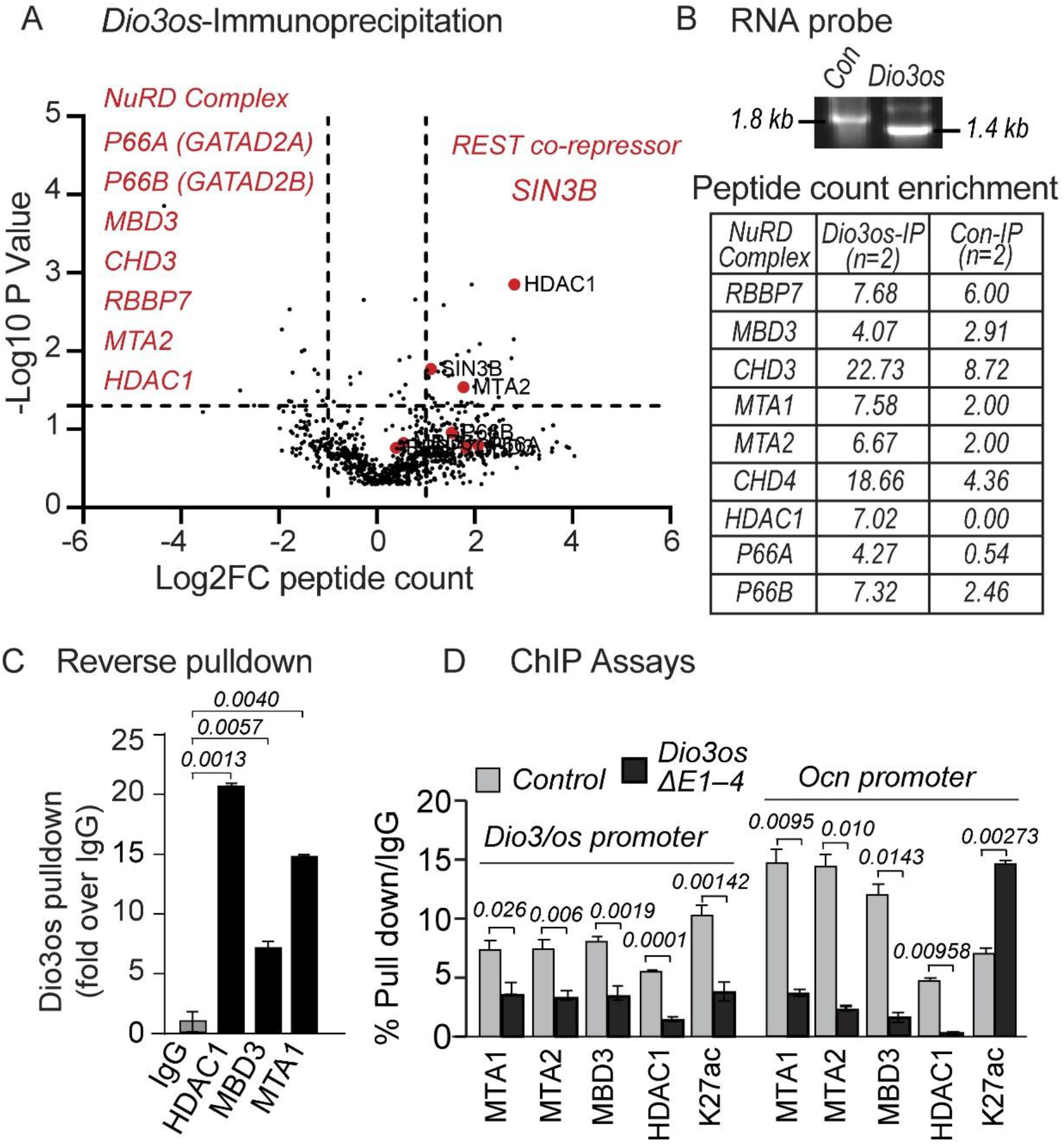
RNA immunoprecipitation with Dio3os, followed by mass spectrometry analysis, identified the HDAC1-guided NuRD complex as part of the Dio3os interactome. (A) Volcano plot from Dio3os RNA immunoprecipitation-mass spectrometry (RIP-MS) showing quantitative protein profiles that are interacting with lncRNA Dio3os (−log10(adjusted P) > 20, log2(FC) > 0.5) between in vitro transcribed, biotin-labeled control luciferase and Dio3os RNAs. NuRD complex members are presented in red. The Volcano plot was generated using the average peptide spectral counts obtained from two independent RIP–MS experiments, with differential enrichment calculated relative to negative-control pulldowns. (B) The 1.8 kb control and 1.4 kb Dio3os transcripts were used in RNA-IP. Average peptide counts from two RNA-IP–MS replicates show enrichment of NuRD components, including HDAC1, RBBP7, MBD3, CHD3, CHD4, MTA1, MTA2, P66A, and P66B. RNA gel shows the biotin-labeled control and Dio3os-specific RNA probe. (C) Reverse Dio3os pulldown with HDAC1, MBD3, and CHD3 antibodies using UV- crosslinked MC3T3-E1 sonicated whole cell lysate. Y-Axis represents fold enrichment over non-specific IgG pulldown. (D) ChIP assays with indicated antibodies against MTA1, MTA2, MBD3, HDAC1, and H3K27ac to determine the binding on Dio3os and Ocn proximal promoters in control (+non-specific sgRNA) and Dio3os(+Dio3os specific sgRNA) CRISPR/Cas9 knockout MC3T3-E1 cells. We performed two independent biological replicates of Dio3os, and control RNA immunoprecipitation-mass spectrometry (RNA-IP-MS) was used to analyze peptide enrichment. For reverse pulldown and ChIP assays, we performed three independent experimental replicates (n = 3) to confirm and validate the reproducibility of the Dio3os interactome. P values are indicated as: NS (P > 0.05), * (P ≤ 0.05), ** (P ≤ 0.01), *** (P ≤ 0.001), and **** (P ≤ 0.0001).

To test whether *Dio3os* recruits NuRD to chromatin, we performed ChIP in WT and Dio3os knockout (ΔE1–4) MC3T3-E1 cells at the *Dio3–Dio3os* and *Ocn* promoters. ΔE1–4 reduced MTA1, MTA2, MBD3, and HDAC1 occupancy at both promoters (Fig. 6D). Strikingly, *Dio3os* deletion produced opposing H3K27ac outcomes, decreased acetylation at the *Dio3–Dio3os* promoter and increased acetylation at the *Ocn* promoter (Fig. 6D). This is the central mechanistic observation of the study. We discovered that the Dio3os–NuRD complex yields distinct chromatin-remodeling and modification outcomes at the loci it activates compared to those it represses. While the molecular basis of this acetylation switch remains to be fully defined, the phenomenon demonstrates a principle of locus-context-dependent NuRD output that may extend to other lncRNA-recruited chromatin platforms.

### Osteoblast-specific CRISPRi and CRISPRa establish bidirectional in vivo control of bone mass

To investigate the *in vivo* function of *Dio3os*, we generated a mature osteoblast-targeted CRISPR interference (CRISPRi) mouse model. This system utilizes a *Col1a1* promoter-driven, tamoxifen-inducible Cre recombinase (*Col1a1-CreERT2*) to conditionally activate dCas9–KRAB expression, alongside two single-guide RNAs (sgRNAs) targeting the regions flanking exon 1 of *Dio3os*. RT-qPCR confirmed 70–80% reduction of *Dio3os* and *Dio3* in femoral bone after induction in both sexes (Fig. 7A). 4-OH-tamoxifen alone had no effect on BV/TV, confirming phenotype specificity (Fig. 7B). μCT analysis at two months post-induction showed increased BV/TV, trabecular number (Tb.N), and trabecular thickness (Tb.Th) with reduced trabecular spacing (Tb.Sp) in both female and male mice (Fig. 7C, 7D, 7E). In females, BV/TV rose from 0.069 to 0.089 (P < 0.001); in males, BV/TV rose from 0.088 to 0.20 (P < 0.05). Histomorphometry confirmed increased osteoblast density (N.ob/BS) and reduced osteoclast density (N.oc/BS), demonstrating a shifted remodeling balance that favors accelerated bone formation over resorption (Fig. 7F). The reciprocal gain-of-function experiment, using osteoblast-targeted CRISPRa (dCas9–SPH) in identical Cre-inducible mice, produced the opposite phenotype: reduced BV/TV, Tb.N, and Tb.Th with increased Tb.Sp; reduced osteoblast and increased osteoclast density (fig. S6A–F). These opposing gain- and loss-of-function phenotypes, observed in both sexes with appropriate controls, establish causal control of bone mass by osteoblast *Dio3os* and rule out off-target or sex-hormone-dependent explanations.

**Figure 7:**
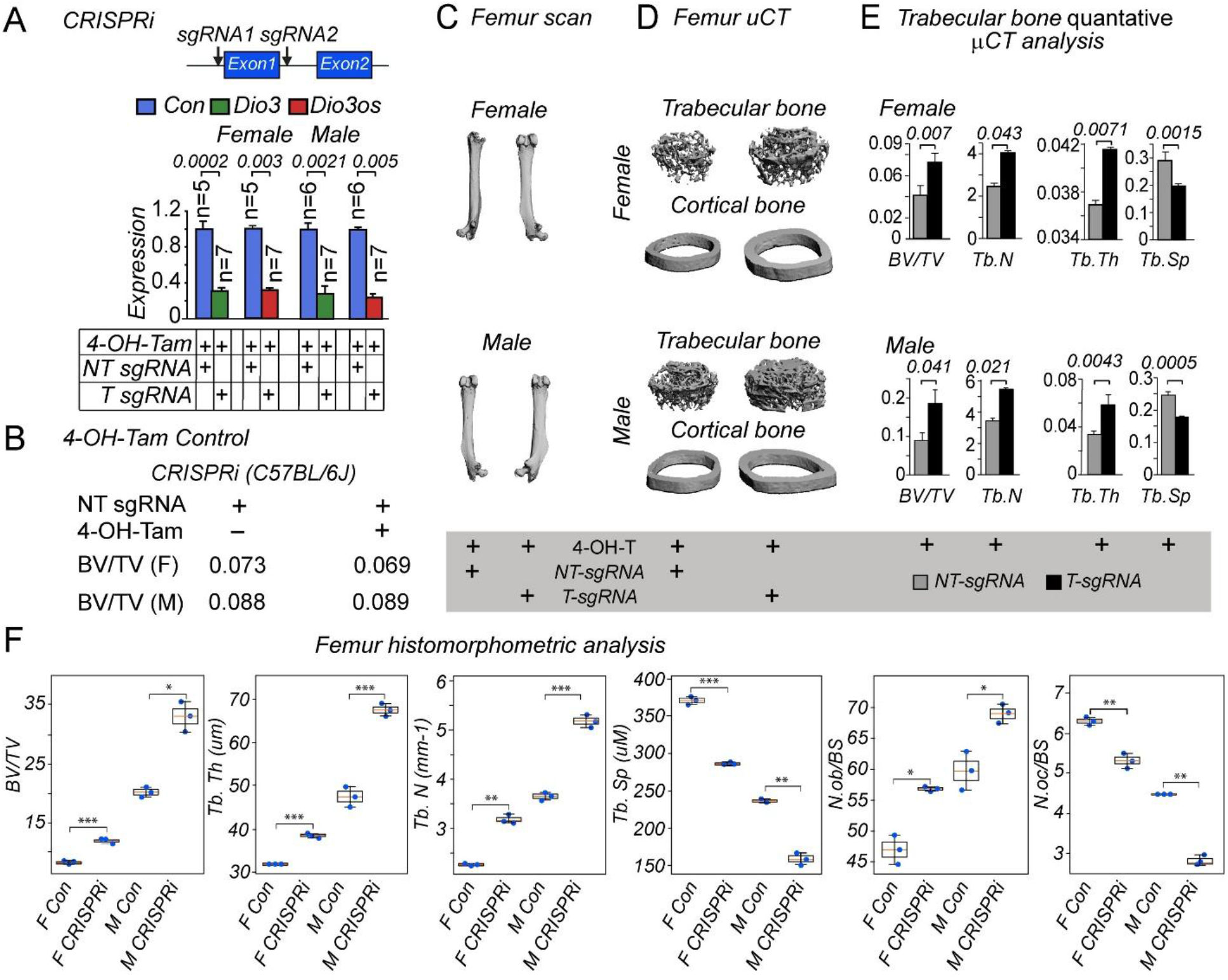
Phenotypic characterization of OB-specific CRISPRi mice to study bone formation (2 months). (A, upper panel) Location of the CRISPRi sgRNAs. (A lower panel) Dio3 and Dio3os mRNA quantification 4 weeks post-injection in femur bone lysates isolated from mice injected with 4-OH- Tamoxifen +/- non-targeting or targeting sgRNAs. CRISPRi groups: Non-targeting sgRNA (n=5 female & n=6 male per group). Targeting sgRNA (n=7 female & n=7 male per group). (B) Femur bone BV/TV ratio from uCT values of CRISPRi female (F) and male (M) 2-month-old mice +/- 4-OH-Tamoxifen. (C upper and lower panels, female & male) Longitudinal microCT scans from CRISPRi + non-targeting sgRNA (NT-sgRNA) (left) and targeting sgRNA (T-sgRNA) (right) mice. Femoral trabecular and cortical microCT scans of two-month-old female (D upper panel) and male (D lower panel) from CRISPRi + NT-sgRNA (left) and T-sgRNA (right) mice. MicroCT quantitation of femoral trabecular bone of female (E upper panel) and male (E lower panel) mice from CRISPRi + NT-sgRNA (grey bar) and T-sgRNA (black bar). Representative graphs illustrate bone volume in relation to tissue volume, trabecular number, thickness, and spacing. Mice were sacrificed at 2 months of age after receiving 10 mg/kg/day of 4-OH-TAM (I/P) and sgRNA lentivirus (109 virus particles/kg/day, I/V) for four consecutive days in one month. Quantitation data here comply with CRISPRi, with n = 7 (T-sgRNA) and n = 5 (NT-sgRNA) female mice and n = 7 (T-sgRNA) and n = 6 (NT-sgRNA) male mice. P values are indicated as: NS (P > 0.05), * (P ≤ 0.05), ** (P ≤ 0.01), *** (P ≤ 0.001), and **** (P ≤ 0.0001). 4-OH-Tam: 4-hydroxytamoxifen. NT-sgRNA: non-targeting sgRNA, and T-sgRNA: targeting sgRNA. (F) CRISPRi (loss-of-function) femur bone histomorphometry analysis showing increased bone formation in female and male mice. Plastic sections of femurs from 8-week-old Control (non-targeting sgRNA + Tamoxifen, n=3 female & n=3 male per group) and CRISPRi (targeting sgRNA + Tamoxifen, n=3 female & n=3 male per group) mice were analyzed for BV (bone volume) /TV (total volume); Tb. N, trabecular number; Tb.Th, trabecular thickness; and Tb. Sp, trabecular space. Data are presented as individual data points with box plots and mean ± SEM (n = 3 per group). Statistical significance was determined using Welch’s t-test comparing control and experimental groups within each sex. Significance levels are indicated as follows: p < 0.05, ** p < 0.01, *** p < 0.001.

## DISCUSSION

This study defines a mode of lncRNA function in which a single transcript uses one chromatin-scaffolding complex to produce two opposite transcriptional outputs at distinct genomic addresses/loci. *Dio3os* recruits the NuRD–HDAC1 complex to its own bidirectional promoter, where it cooperates with HIF1α to activate *Dio3* in cis. In trans, the same *Dio3os*–NuRD complex represses thyroid-hormone–responsive osteogenic gene promoters.

ChIP analysis of the *Dio3–Dio3os* and *Ocn* loci reveals that *Dio3os* deletion reduces NuRD occupancy at both sites but produces opposing H3K27 acetylation outcomes decreased at the activating promoter, increased at the repressed promoter. This locus-context-dependent NuRD output is, to our knowledge, the first demonstration of bidirectional cis/trans regulation by a mammalian lncRNA scaffold and provides a framework for understanding how the same chromatin remodelers can support both activation and repression.

Several features distinguish this mechanism from established lncRNA models. First, the cis activity is mediated through a defined RNA element (exons 2–4) that is dispensable for *Dio3os* trans repression of distal osteogenic targets establishing structure–function resolution at the level of individual RNA exons, which is rarely achieved for mammalian lncRNAs. Second, the integration of chromatin regulation with intracellular hormone metabolism through a single regulatory unit *Dio3os* activates *Dio3* (which inactivates T3/T4 to rT3/T2) while simultaneously suppressing TH-responsive osteogenic transcription couples two regulatory layers that have historically been studied independently. Third, the bidirectional in vivo causality, established with reciprocal CRISPRi (loss-of-function) and CRISPRa (gain-of-function) in both sexes, provides genetic evidence for the model at organism scale.

Our data support a working model in which *Dio3os* scaffolds NuRD–MTA1–HDAC1 to deacetylate HIF1α, stabilizing it and increasing its transcriptional activity at the *Dio3–Dio3os* promoter (^43–45^). The same complex, in cooperation with rT3 produced by *Dio3*, modulates chromatin accessibility at TH-responsive osteogenic genes. RUNX2, which binds the bidirectional promoter and represses its activity at late differentiation, may itself undergo deacetylation by HDAC1, contributing to context-dependent regulation. Although class I HDAC inhibition globally promotes bone formation, our findings reveal a locus-specific role of HDAC1 in *Dio3os*-mediated chromatin remodeling that operates in the opposite direction at *Dio3* versus osteogenic genes emphasizing that HDAC1 function is shaped by its scaffolding context, not its global enzymatic activity.

The findings have implications beyond bone biology. Imprinted lncRNAs at endocrine-relevant loci have been associated with multiple cancers (^29–31^), and the *Dio3os*–NuRD–HIF1α axis we define here may operate in other contexts where hypoxia-responsive chromatin and hormone signaling converge. The principle that a single lncRNA can use one chromatin complex to produce opposite outputs at distinct loci is mechanistically generalizable to other NuRD-, PRC2-, and HDAC-recruiting RNAs (^46–49^), and provides a conceptual framework for the long-standing question of how broad-specificity chromatin remodelers achieve gene-selective regulation. Clinically, our work positions *Dio3os* as a chromatin-level regulator of skeletal homeostasis. Consumptive hypothyroidism, a rare pediatric condition in which aberrant DIO3 inactivates thyroid hormone and impairs skeletal development, emerges as a candidate setting in which *Dio3os*-targeted approaches might restore TH balance and bone formation. More broadly, the in vivo phenotype demonstrates that modulating an imprinted lncRNA can produce robust skeletal outcomes in adult mice without obvious off-target effects, supporting the feasibility of lncRNA-directed therapeutic strategies for endocrine-responsive bone disease.

Several open questions remain. The molecular basis of the context-dependent H3K27ac switch why NuRD recruitment produces increased acetylation at *Dio3–Dio3os* but decreased acetylation at *Ocn* requires structural and biochemical follow-up, possibly involving locus-specific cofactors that gate NuRD’s HDAC versus chromatin-remodeling activities. The mechanisms by which rT3 and T2 stimulate the bidirectional promoter, which appear distinct from T3-driven regulation, warrant additional investigation. Finally, the conservation of this bifunctional architecture in human *DIO3OS* variants and in non-osteogenic tissues will determine its relevance to thyroid-hormone–related disease beyond bone.

## Materials and Methods

### Cell culture

HEK293T (# ATCC CRL-3216, human kidney) and MC3T3-E1 (#CRL-2593, murine calvarial preosteoblast) cell lines were obtained from the American Type Culture Collection (ATCC, Manassas, VA). HEK293T cells were cultured in DMEM, while MC3T3-E1 cells were maintained in αMEM (#12491015 and #12561056, Invitrogen/Gibco, Carlsbad, CA). The culture media were supplemented with 10% fetal bovine serum (FBS; Atlanta Biologicals, Lawrenceville, GA), 2 mM L-glutamine (Thermo Fisher Scientific, Waltham, MA), 50 µg/ml 100 units/ml penicillin, and 100 μg/ml streptomycin (Thermo Fisher Scientific, Waltham, MA). Osteogenic differentiation was induced by adding ascorbic acid and 3–5 mM β-glycerophosphate (Sigma-Aldrich, Cambridge, MA) to the growth medium. Medium was replaced every 2 days throughout all experiments. All cells were maintained at 37 °C in a humidified atmosphere containing 5% CO2.

### LncRNA-Dio3os knockout and polyadenylation insertion using CRISPR/CAS9

To knock out specific genomic sequences of lncRNA-Dio3os, we designed a pair of single guide RNAs (sgRNAs) targeting sequences flanking the region of interest. The sgRNAs were designed using Benchling.com and cloned into the Lenti CRISPR v2 vector (Addgene #52961, Watertown, MA).

Two sets of sgRNAs were created:

gRNA-Dio3os ΔE1 Forward and gRNA-Dio3os ΔE1 Reverse were designed to target the 5’ and 3’ intronic sequences flanking exon 1 of Dio3os.

gRNA- Dio3osΔE1-4 Forward and gRNA- Dio3osΔE1-4 Reverse were designed to target the 5’ intronic sequence flanking exon 1 and the 3’ flanking sequence of exon 4.

The optimal gRNAs were selected from Benchling.com and cloned into the lentiCRISPRv2 vector. The lentiCRISPRv2-Dio3os sgRNA constructs were co-transfected with the pMD2.G (Addgene Plasmid #12259, Watertown, MA) and pCMV delta R8.2 (Addgene, Plasmid #12263, Watertown, MA) viral packaging vectors into HEK-293T cells. The viral supernatants were collected 48 hours of post-transfection. MC3T3-E1 cells were infected with viral particles at 60–70% confluence for 48 hours and subsequently selected with puromycin. Monoclonal cell lines were isolated and confirmed by PCR and sequencing with primers flanking the sgRNA.

sgRNA information was listed as follows:

To delete Dio3os exon 1 (ΔE1), the following sgRNAs were used.

sgRNA #1: CGGACGTTGCTCTCTGCCCC

sgRNA #2: GACGTTGCTCTCTGCCCCGG

sgRNA #3: TCAACAGGGAACGTGTGCTG

sgRNA #4: TCCTAATCTGTCTCTGCGCG

sgRNA #5: TGGCGCACTGCTGGAGACTT

To delete Dio3os Exon1-4 (ΔE1-4), the following sgRNAs were used.

sgRNA #1: CGGACGTTGCTCTCTGCCCC

sgRNA #2: GACGTTGCTCTCTGCCCCGG

sgRNA #3: CTGAAAGAGGTGCTTAGCCC

sgRNA #4: AGGTCATCTAGGACTGGCTCCG

sgRNA #5: GTTCTGCCCTCTGCACACGG

sgRNA #6: CTTACTGAAAACCGGGTTTG

We successfully generated sgRNAs targeting exon 1 and exons 1-4 and verified the Dio3os CRISPR-Cas9 exon knockouts (ΔE1 and ΔE1–4) through PCR genotyping using allele-specific primers, Sanger sequencing of the junctions, and genomic BLAST analysis within the Dio3os locus without affecting the Dio3 coding region (**fig. S3 A–G**). We used pooled sgRNAs to knock out Dio3os exon 1 or exons 1–4. Hence, the double-strand break (DSB) positions and repair junctions varied slightly due to the randomly determined nature of non-homologous end joining (NHEJ). As a result, we observed sequence differences in the residual exon 1 region, which are typical of CRISPR-mediated deletions and do not impact functional knockout efficiency (^50,51^). To insert the pAS (polyadenylation) sequence into the Dio3os genomic locus, we used a donor plasmid kindly received from Dr. Eric S. Lander’s lab (https://www.broadinstitute.org/lander-lab). The 3’ homologous arm of Dio3os was cloned into the donor vector using the restriction enzyme ClaI. In comparison, the 5’ homologous arm was cloned using the restriction enzyme AflII, following the protocol described by (^52^). Briefly, the donor plasmid was packaged into a virus along with the CRISPR/Cas9 plasmid, followed by selection as described above. Monoclonal cell lines were then isolated and confirmed by PCR and sequencing.

5HA exon 1-2 FW: CCTTAGTACCCTTAAGCTAGGGGTAGCTGTTGCCTGAAC

5HA exon 1-2 RV: CAACGTTGCCCTTAAGTCTCCAGCAGTGCGCCAC

3HA exon 1-2 FW: CTAGTACGCGTATCGATCTTGGGTTCAGGGGCGG

3HA exon 1-2 RV: CCCTCGACGGTATCGATGCCAGAGTCTCTTCTCACtttaccatct

5HA exon2-3 FW: CCTTAGTACCCTTAAGCGAACTCGGAGCGCGGC

5HA exon2-3 RV: CAACGTTGCCCTTAAGGGCTACTTGAGGTCGACTtttccggac

3HA exon2-3 FW: CTAGTACGCGTATCGATCAGAGGGCGCCCAGGTAG

3HA exon2-3 RV: CCCTCGACGGTATCGATGCCAGAGTCTCTTCTCACtttaccatc

5HA_exon 3-4 FW: CCTTAGTACCCTTAAGCAGGCCTTCGACTTCCCTg

5HA_exon 3-4 RV: CAACGTTGCCCTTAAGCCAGGGCATGATGAGATGgg

3HA_exon 3-4 FW: CTAGTACGCGTATCGATTTACATGGGCTCAGTCTTCcttg

3HA_exon 3-4 RV: CCCTCGACGGTATCGATGGCCCCTCCCAGAAGTC

sgRNA sequence for pAS into Dio3os Exon1-2

sgRNA #1 AAGTCTCCAGCAGTGCGCCA;

sgRNA #2: CGCGCAGAGACAGATTAGGA

sgRNA sequence for pAS into Dio3os Exon 2-3

sgRNA #1 CTGGGCTACTTGAGGTCGAC

sgRNA #2: TCTGGGCTACTTGAGGTCGA

sgRNA sequence for pAS into Dio3os Exon 3-4:

sgRNA #1 AGACTGAGCCCATGTAACCA

sgRNA #2: TGTAACCAGGGCATGATGAG

We performed three independent experimental replicates (n = 3) of Dio3os exons CRISPR knockout and exons polyadenylation to confirm and validate the reproducibility of Dio3os expression.

### Construction of lncRNA-Dio3os overexpression plasmid and stable overexpression in MC3T3-E1 cells

To construct the lncRNA-Dio3os overexpression plasmid, lncRNA Dio3os was amplified using the CloneAmp™ HiFi PCR System (639298, Takara Bio, San Jose, CA). The primers used were: Forward (FW): 5’ GTATTCTAGAGCTAGGAGACTGGAGCGCCCGAA 3’; Reverse (RV): 5” ATTCGAATTCGCTAGTTCTCATTGCACAAAGCTTTATTGGGCC 3’ The amplified product was cloned into the PCDH-Mscv-coGFP (System Biosciences, SBI, Palo Alto, CA) vector at the Nhe I restriction enzyme site. The correct sequence of the construct was confirmed through Sanger sequencing at the UAB Sequencing Core Facility (Genomic Core, UAB, Birmingham, AL). The PCDH-LncRNA Dio3os plasmid was co-transfected with the pMD2.G and pCMV delta R8.2 packaging plasmids (Addgene, Watertown, MA) into HEK293T cells to produce lentivirus. The lentiviral supernatant was collected and used to infect MC3T3-E1 cells. GFP-positive cells, indicating successful infection and expression, were sorted using fluorescence-activated cell sorting (FACS).

### RNA Isolation and Quantitative Real-Time PCR

Total RNA was isolated using TRIzol® Reagent (Ambion/Life Technologies, Carlsbad, CA), followed by phenol-chloroform extraction. Complementary DNA (cDNA) was synthesized through a reverse transcription reaction using random hexamer and oligo-dT primers using Advantage® RT-for-PCR Kit (639506, Takara Bio, San Jose, CA). Quantitative real-time PCR (qPCR) was performed using Luna Universal qPCR Master Mix (New England Biolabs, MA) and gene-specific primers. The reactions were run on a 7500 Fast Real-Time PCR System (Applied Biosystems, Waltham, MA).

RT-qPCR Primers were listed as follows:

Dio3os: Forward: 5’ CGGGAGAGACCAGCTACCTA 3’

Reverse: 5’ GCCACTTTCACAGGCTGTTC 3’

Dio3: Forward: 5’ CCCATATGCGTATCAGACGA 3’

Reverse: 5’ CCTTGTGCGTAGTCGAGGAT 3’

Runx2: Forward: 5’ CGGCCCTCCCTGAACTCT 3’

Reverse: 5’ TGCCTGCCTGGGATCTGTA 3’

Ocn: Forward: 5’ CTGACAAAGCCTTCATGTCCAA 3’

Reverse: 5’ GCGCCGGAGTCTGTTCACTA 3’

Alp: Forward: 5’ CCAACTCTTTTGTGCCAGAGA 3’

Reverse: 5’ GGCTACATTGGTGTTGAGCTTTT 3’

GAPDH: Forward: 5’ AGGTCGGTGTGAACGGATTTG 3’

Reverse: 5’ TGTAGA CCATGTAGTTGAGGTCA 3’

### Western Blot Analysis

Cells were lysed in FA buffer (1 mM EDTA, pH 8.0; 50 mM HEPES-KOH, pH 7.5; 140 mM NaCl) supplemented with 1% Triton X-100, 0.1% sodium deoxycholate, 1 mM PMSF, protease inhibitors (Roche), and MG132 (Sigma-Aldrich). Lysates were Dounce-homogenized and sonicated on ice (2% power, 10 s ×3). Equal protein amounts (30 µg) were resolved by SDS–PAGE (10%) and transferred to PVDF membranes. Proteins were detected using the LI-COR Bio-Imaging System (LI-COR Biosciences). Primary antibodies: HDAC1 (Santa Cruz, sc-81598; 1:500); HIF-1α (Novopro, 111345; 1:500); RUNX2 (Santa Cruz, sc-10758; 1:1,000); BRG1 (Santa Cruz, sc-10768; 1:1,000); B-tubulin (loading control, Proteintech, 11224-1-AP, 1:10,000) and GAPDH (loading control, Abcam, mAbcam 9484).

### Human LncProfilers qPCR Assay

As previously described, total RNA was extracted using TRIzol® Reagent and then synthesized into cDNA from differentiated MC3T3-E1 osteoblast cells at the specified time points. The human lncRNA plate Human LncProfilers™ qPCR Array Kits (Cat # RA900A-1) was obtained from System Biosciences (SBI, Palo Alto, CA), and the manufacturer’s protocol was followed. The kits include assays in preformatted plates for well-annotated human lncRNAs, with three endogenous reference RNA controls per plate. All of the lncRNAs on the qPCR array have validated primer sets for well-annotated lncRNAs that are registered in the lncRNA database created by Dr. John Mattick (www.lncrnadb.org). Real-time quantitative PCR (RT-qPCR) was performed in three independent experimental replicates (n = 3) to ensure reproducibility, and the data were analyzed using the ΔΔCt method to calculate fold changes.

### Chromatin Immunoprecipitation Sequencing (ChIP-Seq) and ChIP-qPCR

The ChIP-seq data were obtained with permission from the authors of “Chromatin dynamics regulate mesenchymal stem cell lineage specification and differentiation to osteogenesis” (^53^) and are sourced from GEO (Series GSE76074). Data were analyzed in IGV and Adobe Photoshop using area-under-the-curve measurements. It was normalized by gene length, scaled, and plotted in 3D with OriginPro 2019. ChIP-seq was performed on bone marrow-derived stem cells (BMSCs) undergoing osteogenic differentiation, following the published protocol (^53^). Crosslinked and sheared chromatins were then immunoprecipitated with Runx2 antibody (M-70, Santa Cruz), H3K4me3 (ab8895, Abcam, Cambridge, MA, USA), H3K27me3 (07-449, Millipore, Burlington, MA), H3K27ac (07-360, Millipore, Burlington, MA), H3K36me3 (ab282572, Abcam, Cambridge, MA) or immunoglobulin G (IgG) (12-370, Millipore, Burlington, MA). Protein-DNA complexes were purified with Protein-G Dynabeads (Invitrogen, Waltham, MA), and DNA libraries were sequenced on the Illumina Genome Analyzer II or HiSeq-1500. Base calls and sequence reads were generated using Illumina CASAVA or bcl2fastq software (versions 1.8 and 1.8.4). At each time point, two independent biological replicates of ChIP-Seq libraries and two input libraries derived from sonicated DNA were prepared. Primer sequences for ChIP qPCR were designed and assessed using RT-qPCR. At each time point, we prepared two independent biological replicates of ChIP-Seq libraries, along with two input libraries derived from sonicated DNA.

For chromatin immunoprecipitation followed by quantitative PCR (ChIP–qPCR; Fig. 9D), control and Dio3os CRISPR knockout (DE1–4) MC3T3-E1 cells were differentiated for 7 days. ChIP was performed on day 7 using standard protocols, and immunoprecipitated promoter DNA was quantified by qPCR. Antibodies used were MTA1 (Santa Cruz Biotechnology, sc-373765), MTA2 (sc-5566), MBD3 (sc-166319), HDAC1 (sc-81598), and histone H3 acetyl-Lys27 (H3K27ac; Abcam, ab4729). qPCR was performed using primers targeting the Dio3os and osteocalcin (OCN) promoter regions.

Dio3os promoter primers

Forward: 5′-CGGCGGGGCTGCAGCCGGGGGCC-3′

Reverse: 5′-GCCGGAGCTCCGGTTCGCTTCCCGCGCGCC-3′

OCN promoter primers

Forward: 5′-CGCAATCACCTACCACAGC-3′

Reverse: 5′-CCGCTAGTCTGTGCTCTCTG-3′

### Assay for Transposase Accessible Chromatin (ATAC) Sequencing

Chromatin accessibility profiles at the Dio3-Dio3os gene loci were assayed during MCM3T3-E1 proliferation on Day 0 and the onset of differentiation on Day 7 as described by Buenrostro et al.(^54^) Cells were lysed in ice-cold lysis buffer (10mM Tris-HCl, pH 7.4, 10mM NaCl, 3mM MgCl2, and 0.1% Igepal CA-630). Lysates were incubated with Tn5 transposase (Illumina Tagment DNA Enzyme and Buffer Kit, Illumina, San Diego, CA, USA) for 30 min at 37°C, followed by DNA purification using the MinElute PCR Purification kit (Qiagen, Germantown, MD, USA). Transposed DNA was amplified with barcoded Nextera primers (Illumina, San Diego, CA, USA) using the NEBNext High-Fidelity 2X PCR Master Mix (New England Biolabs, Ipswich, MA, USA). The barcoded DNA libraries were sequenced at GeneWIZ, and Bowtie2 was used to map the reads to the mouse genome (mm10). Sequencing was performed on an Illumina HiSeq platform with 2×150 bp reads, yielding approximately 150 million raw reads per sample. At each time point (days 0 and 7 for Figure 3, and day 14 for Figure 8), we prepared one ATAC-Seq library for each time point, along with the corresponding input libraries, and verified them using three replicate ATAC-qPCR assays performed on DNA treated with Tn5 transposase.

### RNA Sequencing and Bioinformatic Analysis

To identify lncRNA Dio3os–regulated thyroid hormone–responsive and osteoblast-specific genes, high-throughput RNA sequencing (RNA-seq) was performed on MC3T3-E1 osteoblasts from control and Dio3os exon 1–4 knockout (ΔE1–4) lines. Total RNA (0.5–1 μg) was extracted using the RNeasy Mini Kit (Qiagen) and quality-checked by Bioanalyzer 2100 (Agilent Technologies). Ribosomal RNA was removed using an rRNA Removal Kit (Illumina) or poly(A) capture method, followed by RNA fragmentation with RNase III. We used an Illumina cDNA Library Prep Kit and constructed the sequencing libraries according to the manufacturer’s protocols (Novogene, Beijing, China). Random hexamer primers and M-MuLV Reverse Transcriptase were utilized to synthesize first-strand cDNA with MuLV Reverse Transcriptase. The second-strand synthesis was performed using DNA polymerase I and RNase H. cDNA fragments were amplified using high-fidelity DNA polymerase and assessed for library quality before sequencing on an Illumina NovaSeq 6000 platform (paired-end 150 bp). Raw sequencing reads were processed using a standard pipeline. Read quality was assessed using FastQC (^55^), and adapters were trimmed with TrimGalore (^56^). Clean reads were aligned to the mouse reference genome (mm10) using HISAT2. (^57^) Gene-level expression was quantified using FeatureCounts (^58^), and normalized as FPKM values with StringTie (^59^). Differentially expressed genes (DEGs) were identified using DESeq2 (^60^) with a false discovery rate (FDR)–adjusted P value < 0.05. Gene Ontology (GO) enrichment analysis was performed using the clusterProfiler package (^61^) and annotated with the Gene Ontology resource (^62^). Both wild-type and knockout libraries showed high sequencing quality, with total mapping rates >97% and uniquely mapped reads >84% (fig. S3A). Transcriptome correlation (R² = 0.957) confirmed reproducibility across biological replicates (Supplementary Fig. 3B). Coverage plots generated with deepTools revealed a complete loss of Dio3os read density and reduced Dio3 coverage in knockout cells, consistent with shared bidirectional promoter regulation.

### Measurement of Dio3 and Dio3os expression with thyroid hormone metabolites

MC3T3-E1 cells were seeded in 6-well plates and treated with the indicated drugs (T2, rT3, and T3) at 100 and 1000 ng/mL for 48 hours. Total RNA was extracted using TRIzol reagent, and cDNA was synthesized via reverse transcription. Relative mRNA expression levels of the target genes were measured using RT-qPCR and calculated using the ΔΔCt method, with GAPDH serving as an internal control. We performed three independent experimental replicates (n=3) to ensure the reproducibility of Dio3 and Dio3os expression.

### Cloning of the Dio3os/Dio3 promoter in the Luciferase reporter construct

Dio3os/Dio3 promoter was amplified from mouse genomic DNA and cloned into pGL3 basic plasmid using the NheI restriction enzyme. The successful plasmid was sequenced and confirmed. The primers were listed below:

Dio3 promoter FW: TCTTACGCGTGCTAGCGGGCTGCGCCGGGCGAGG

Dio3 promoter RV: TCGAGCCCGGGCTAGCGGTCGAGGGTGGAGCGGT

Dio3os promoter FW: TCTTACGCGTGCTAGCTCGGTGCTCTGGGAGCTC

Dio3os promoter RV: TCGAGCCCGGGCTAGCGGGCTGCGCCGGGCGAGG

### Dio3-Dio3os Promoter Luciferase Reporter Assay

The Dio3/Dio3os luciferase reporter vector was co-transfected with the indicated plasmids (Runx2, Hdac1, Hif1a, and Brg1), or with HDAC inhibitors, thyroid hormone, and its derivatives, into MC3T3-E1 osteoblast cells along with Dio3os (WT) and CRISPR-Cas9–mediated exon-deleted versions of Dio3os stable constructs. Samples were collected 48 hours after transfection to detect luciferase activity. We performed three independent experimental replicates (n = 3) of Dio3-Dio3os promoter-reporter assays to confirm and validate the reproducibility of Dio3-Dio3os bidirectional promoter activity.

### ALP Staining

To detect alkaline phosphatase (ALP) activity, control, and Dio3os CRISPR deleted (ΔE1-4) MC3T3-E1 osteoblast cells at differentiation day 14 were fixed using cold, freshly prepared 10% neutral buffered formalin for 15 minutes. Following fixation, the cells were incubated at room temperature for 2 hours in a solution containing 0.4 mg/ml naphthol AS-MX phosphate (catalog no. N5000, MilliporeSigma, St. Louis, MO) and 0.08 mg/ml Fast Red TR (catalog no. F8764, MilliporeSigma, St. Louis, MO) to facilitate color development. After incubation, the cells were washed with distilled water, air-dried, and photographed.

### Von Kossa Staining

To detect phosphate-rich calcium deposits in control and Dio3os CRISPR knockout (ΔE1-4) MC3T3 cells, we first fixed cells at differentiation day 21 in 4% paraformaldehyde for 10–15 minutes at room temperature, then rinsed them with distilled water. The samples were then incubated with freshly prepared 5% silver nitrate solution for 30–60 minutes under strong light (UV or full-spectrum lamp), which photo-reduces silver ions to metallic silver at sites of phosphate accumulation, marking mineralized areas. After exposure, the samples were thoroughly washed with distilled water and treated with 0.5% sodium thiosulfate for 5 minutes to remove unreacted silver, followed by additional rinses. Samples were then either imaged directly (for cell culture) or dehydrated through graded ethanol, cleared in xylene, and mounted with a coverslip for tissue sections. Mineralized areas appeared black or dark brown. We conducted three independent experimental replicates (n = 3) of ALP and Von Kossa staining using Control and Dio3os CRISPR (ΔE1-4) knockout MC3T3-E1 cells to confirm the reproducibility of the Dio3os knockout effect on osteoblast activity and mineralization.

### Generation of Dio3os tamoxifen-induced conditional CRISPRi, CRISPRa, and control mice

Homozygous CRISPRi (Rosa26-LSL-dCas9-KRAB, Strain # 033066, Jackson laboratory, Bar Harbor, ME) and CRISPRa (pb-CAG-Cas9*,-EGFP, Strain # 031645, Jackson laboratory, Bar Harbor, ME) mice bred with Col1a1-CreERT2 (Col1a1-cre/ERT2, Strain # 016241, Jackson laboratory, Bar Harbor, ME) mice to generate homozygous Dio3-Dio3os CRISPRi and CRISPRa mice that are hemizygous for the Cre transgene. PCRs were performed for genotype confirmation. We evaluated the experimental (CRISPRi and CRISPRa + 4-OH-TAM + target sgRNA) and control groups (CRISPRi and CRISPRa + 4-OH-TAM + non-target sgRNA) to examine the effect of CRISPRi and CRISPRa. The UAB Transgenic & Genetically Engineered Models (TGEM) core helped us to successfully and quickly generate CRISPR models. All mouse experiments were performed following the Institutional Animal Care and Use Committee (IACUC) guidelines at the University of Alabama at Birmingham (Animal Protocol No. 21319).

### Tamoxifen induction, viral sgRNA production, and in vivo delivery

To induce CreERT2-mediated recombination in osteoblasts, 1-month-old mice were administered 4-OH-tamoxifen (10 mg/kg/day, intraperitoneal) for 4 consecutive days. U6-driven lentiviral sgRNA vectors (lenti U6-sgRNA/EF1α-mCherry; Addgene #114199, Watertown, MA) were produced by co-transfecting 293T cells with psPAX2 (Addgene #12260, Watertown, MA) and pMD2.G (Plasmid #12259, Addgene, Watertown, MA) plasmids, and viral supernatants were collected at 48 and 72 h. Parallel to tamoxifen induction, mice received systemic delivery of lentiviral sgRNAs (targeting or non-targeting) via tail vein injection at a dose of 1×10⁹ viral particles/kg, administered every 3–4 days for one month.

### Micro-CT analysis of bone parameters

At the end of the one-month induction period (2 months of age), the mice were euthanized, and their femurs were harvested for quantitative microCT analysis. Scans were acquired at a 10 μm voxel size, and 25 longitudinal slices per femur were reconstructed. For trabecular assessment, 100 slices immediately distal to the growth plate were analyzed, and 50 slices from the mid-diaphyseal region were used for cortical measurements. Specific microCT parameters (sigma = 1.2, support = 2, threshold = 180) were applied to quantify bone architecture, including trabecular bone volume fraction (BV/TV), trabecular thickness (Tb. Th), trabecular number (Tb.N), and trabecular separation (Tb.Sp) in the proximal and distal femoral metaphysis.

### Bone Histomorphometry

Femora from 8-week-old littermates were harvested, fixed in 4% paraformaldehyde, and processed for histological analysis. For structural histomorphometry, bones were dehydrated in graded ethanol and embedded in methyl methacrylate without decalcification. Longitudinal sections (5–7 μm) were prepared from the distal femoral metaphysis and stained with Goldner’s trichrome. Static histomorphometric parameters, including bone volume per tissue volume (BV/TV), trabecular thickness (Tb.Th), trabecular number (Tb.N), and trabecular space (Tb.Sp), were quantified using OsteoMeasure software (OsteoMetrics, Decatur, GA) within a defined region of interest located 0.5–2.0 mm proximal to the growth plate. Osteoblast number per bone surface (Ob.N/BS) was determined by identifying cuboidal osteoblasts lining the trabecular bone surface in Goldner’s trichrome-stained sections. For osteoclast analysis, decalcified sections of paraffin-embedded bones were stained for tartrate-resistant acid phosphatase (TRAP). Osteoclast number per bone surface (Oc.N/BS) was quantified by counting TRAP-positive multinucleated cells along the trabecular surface. All histomorphometric measurements were performed in a blinded manner in accordance with the standardized nomenclature established by the American Society for Bone and Mineral Research (ASBMR).

### dCAS9-KRAB Mice Genotype

Forward Primer: 5’ GCAGCCTCTGTTCCACATACAC 3’

Reverse Primer 1: 5’ TAAGCCTGCCCAGAAGACTC 3’

Reverse Primer 2: 5’ AAAGTCGCTCTGAGTTGTTAT 3’

Products: WT= 235bp; Mut=162bp

dCAS9-SPH Mice Genotype:

Forward Primer: 5’ CACCATCTCCCTGCTGACA 3’

Reverse Primer: 5’ CTGAACTTGTGGCCGTTTAC 3’

Products: Homozygous = 264 bp

Col1a1-Cre ERT2 (2.3kb) Mice Genotype:

Forward Primer: 5’ ACAATCAAGGGTCCCCAAAC 3’

Reverse Primer: 5’ CCAGCCGCAAAGAGTCTACA 3’

Products: Hemizygous = 118 bp

sgRNA sequence: The following sgRANAs are cloned into the lenti U6-sgRNA/EF1α-mCherry vector.

CRISPR Inhibition-dCas9-KRAB: sgRNA1/sgRNA2

CRISPR Activation-dCas9-SPH: sgRNA1/sgRNA3

sgRNA1: 5’ CTCCAGTCTCGACGTTCCCG 3’

sgRNA 2: 5’ ACGCGCAGAGACAGATTAGG 3’

sgRNA 3: 5’ GCGAGGCAAGCGGCGAGAG 3’

### Biotin-labeled Dio3os RNA Pulldown and Mass Spectrometry Analysis

The lncRNA Dio3os sequence was cloned into the pET28b (+) vector and transcribed in vitro using the Biotin-16-UTP RNA Labeling Kit (Roche, Indianapolis, IN) to generate biotin-labeled Dio3os RNA. Nuclear extracts from MC3T3-E1 cells were incubated with biotinylated Dio3os RNA overnight at 4 °C, followed by pulldown using streptavidin-coated magnetic beads (Thermo Fisher Scientific, Waltham, MA). Beads were washed five times with Buffer A (150 mM KCl, 15 mM Tris, pH 7.4, 5 mM EDTA, 0.5 mM DTT, 0.5% NP-40, 0.5% sodium deoxycholate, RNaseOUT, and protease inhibitors including Halt PMSF and MG132). Bound proteins were eluted in 1× LDS sample buffer at 96 °C for 10 min, reduced and denatured at 70 °C for 10 min, and separated on a 10% Bis-Tris gel. Gels were stained overnight with colloidal Coomassie and submitted to the UAB LC–MS Core Facility for mass spectrometry. Mass spectrometry data were analyzed using the SEQUEST algorithm (Proteome Discoverer v1.4) with high-resolution Orbitrap MS2 datasets, as described previously (^63,64^). Quantitative proteomics data were processed by removing outliers (ROUT test, Q = 1%) and evaluated by two-way ANOVA with Tukey’s post hoc test and Benjamini–Hochberg FDR correction (q = 5%). Functional annotation used SAINT, UniProt, Protein Atlas, and subcellular localization databases. Statistical analysis and graphical presentation were performed in GraphPad Prism v9.2.0. Spectral counts were normalized, and relative protein abundance was expressed as a percentage of total adjusted spectral counts. Two independent experiments were combined, and mean Dio3os-interactome values were normalized to negative control (NC) pulldowns to calculate log₂ fold changes.

### Reverse RNA Pulldown Assay

MC3T3-E1 cells were cultured overnight and UV-crosslinked to stabilize RNA–protein interactions. Cells were irradiated at 4,000 × 100 µJ/cm² for 30 s, washed with ice-cold 1× PBS, and irradiated again at 2,000 × 100 µJ/cm² for 30 s. Cells were harvested and pelleted at 5,000 × g for 5 min at 4 °C. Approximately 1 × 10⁷ cells were lysed in PXL buffer (0.1% SDS, 0.5% NP-40, 0.5% sodium deoxycholate, 1 mM DTT in 1× PBS) supplemented with RNaseOUT and incubated on ice for 10 min. Lysates were sonicated (30 s ON / 1 min OFF, three cycles), treated with DNase I at 37 °C for 15 min, and clarified by centrifugation at 13,000 rpm for 30 min at 4 °C. Ten percent of the supernatant was reserved as input.

The remaining lysate was incubated overnight at 4 °C with the indicated antibodies, followed by three washes with PXL buffer. RNA was extracted using TRIzol, and Dio3os enrichment was quantified by qPCR using gene-specific primers. Relative enrichment was calculated as log₂ fold change and normalized to the IgG control. Antibodies: MBD3 (Santa Cruz Biotechnology, sc-166319); MTA1 (Santa Cruz Biotechnology, sc-373765); HDAC1 (Santa Cruz Biotechnology, sc-81598)

**qPCR primers:** Dio3os forward: 5′-CGGGAGAGACCAGCTACCTA-3′, Dio3os reverse: 5′-GCCACTTTCACAGGCTGTTC-3′

### Statistical analysis

All measured data are presented as the mean ± standard deviation (SD), and Welch’s t-test was used to compare groups using GraphPad Prism8. P-value < 0.05 was considered statistically significa nt. P values are indicated as: NS (P > 0.05), * (P ≤ 0.05), ** (P ≤ 0.01), *** (P ≤ 0.001), and **** (P ≤ 0.0001).

## Acknowledgments

We want to express our gratitude to the members of the Department of Oral and Maxillofacial Surgery and the Institute of Oral Health Research at the School of Dentistry, University of Alabama at Birmingham (UAB) for their support and insightful discussions. Special thanks to Solo Jacqueline Bergill-Gentile for her feedback and encouragement.

We also appreciate Dr. Gary Stein, Dr. Janet Stein, and Dr. Jane Lian from the University of Vermont for providing BMSC ChIP-sequencing data, as well as the UAB Heflin Center for Genomic Science Core for assistance with sequencing analyses, and the UAB Animal Resources Program (ARP) for caring for the CRISPRi and CRISPRa mouse models.

## Funding

This work was funded by the National Institute of Arthritis and Musculoskeletal and Skin Diseases (NIAMS) under grant R01AR069578. The authors are solely responsible for the content, which does not necessarily represent the official views of the NIH.

## Author Contributions

Y.C. is responsible for data curation, validation, investigation, visualization, methodology, and writing (review and editing). B.W. focused on conceptualization, data curation, formal analysis, and methodology. M.R. contributed to data curation, experimental work, data analysis, and methodology. S.M. engaged in conceptualization, data organization, and writing (review and editing). S.K. contributed to data organization, experimental execution, and manuscript review and editing. L.H.A. contributed to experimental conceptualization and design, and manuscript review and editing. Q.H. was involved in conceptualization, resource management, formal analysis, supervision, funding acquisition, validation, investigation, visualization, figure preparation and editing, methodology, writing (original draft), writing (review and editing), and finalizing the manuscript.

## Competing interests

The authors declare no competing interests.

## AI-assisted language editing

AI-assisted language editing was accomplished using ChatGPT to enhance clarity and readability. No scientific content, data, or interpretations were generated by the AI, and all material was reviewed and approved by the authors.

## Data, code, and materials availability

The RNA-seq and ATAC-seq datasets generated in this study have been deposited in the NCBI Gene Expression Omnibus (GEO) under accession numbers GSE300822 (RNA-seq) and GSE300729 (ATAC-seq). The mass spectrometry proteomics data have been deposited under Project accession PXD065537. Previously published ChIP-seq datasets used for comparative analysis were obtained from the GEO database under accession number GSE76074. All plasmids, CRISPR sgRNA sequences, primers, and reagents used in this study are described in the Methods section and are available from the corresponding author upon reasonable request. All source data underlying the figures are provided as Source Data files.

(1) To review GEO accession GSE300822 (RNA-seq):

Go to:

https://nam12.safelinks.protection.outlook.com/?url=https%3A%2F%2Fwww.ncbi.nlm.nih.gov%2Fgeo%2Fquery%2Facc.cgi%3Facc%3DGSE300822&data=05%7C02%7Cyuechuan%40uab.edu%7C70421c19881145a4d3de08ddb40bfe50%7Cd8999fe476af40b3b4351d8977abc08c%7C1%7C0%7C638864685805871674%7CUnknown%7CTWFpbGZsb3d8eyJFbXB0eU1hcGkiOnRydWUsIlYiOiIwLjAuMDAwMCIsIlAiOiJXaW4zMiIsIkFOIjoiTWFpbCIsIldUIjoyfQ%3D%3D%7C0%7C%7C%7C&sdata=8RB%2Bss3Nv36d8%2FEtfg74%2FyJNLxs0qSAqBH50bT%2F4DFM%3D&reserved=0

Enter token mpgvausyjzsbhkj into the box.

(2) To review GEO accession GSE300729 (ATAC-seq)

Go to:

https://nam12.safelinks.protection.outlook.com/?url=https%3A%2F%2Fwww.ncbi.nlm.nih.gov%2Fgeo%2Fquery%2Facc.cgi%3Facc%3DGSE300729&data=05%7C02%7Cyuechuan%40uab.edu%7Cd865652773d14cd3654108ddb40bedc2%7Cd8999fe476af40b3b4351d8977abc08c%7C1%7C0%7C638864685514309166%7CUnknown%7CTWFpbGZsb3d8eyJFbXB0eU1hcGkiOnRydWUsIlYiOiIwLjAuMDAwMCIsIlAiOiJXaW4zMiIsIkFOIjoiTWFpbCIsIldUIjoyfQ%3D%3D%7C0%7C%7C%7C&sdata=Xnis%2F5CMmR2V0O9l76fI6Z%2Bf%2BP%2FrJuTbm6eXNDiDktw%3D&reserved=0

Enter token knwneyosfnmnjkb into the box.

(3) The mass spectrometry proteomics data:

Project accession: PXD065537 Token: HhTTY3BwL2Og

## Notes

### Competing Interest Statement

The authors have declared no competing interest.

### Summary of Updates

Dear bioRxiv Editorial Team, I hope you are doing well. I am writing on behalf of all co-authors to request an update to the author list for our bioRxiv preprint. Title: LncRNA Dio3os regulates neighboring gene Dio3 and impacts osteogenesis in trans Preprint DOI: 10.1101/2025.04.14.647401 During the continued development and revision of this manuscript, we performed additional experiments, incorporated new data, expanded the analyses, and substantially revised the manuscript. As part of these efforts, the following researchers made significant scientific contributions that meet the criteria for authorship: Shazia Khan Lubana H. Afreen Old Author list: Yuechuan Chen, Benjamin Wildman, Mohammad Rehan, Snehasis Mishra, Quamarul Hassan New Author list: Yuechuan Chen, Benjamin Wildman, Mohammad Rehan, Snehasis Mishra, Shazia Khan, Lubana H. Afreen, Quamarul Hassan Their contributions included participating in the generation and validation of new experimental data, assisting with data analysis and interpretation, contributing to the revision and improvement of the manuscript, and providing important intellectual input during the preparation of the revised version. Accordingly, we respectfully request that the author list be updated to include these two authors. All authors have reviewed and approved this request and agree with the revised authorship. We are happy to provide any additional information or written confirmation from all co-authors if required. Thank you very much for your time and consideration. We appreciate your assistance and look forward to your guidance regarding the authorship update. Sincerely,

