## Supplemental Information for "A bifunctional imprinted lncRNA scaffolds NuRD–HDAC1 to couple cis-activation of Dio3 with trans-repression of osteoblast chromatin"

**Supplemental Materials and Methods**

**miRNA-1247 Knockdown Assay**

MC3T3-E1 cells were seeded in a 6-well plate at approximately 50% confluence and transfected with control and miR-1247 specific synthetic small interfering RNA at a concentration of 100 nM for 2 days. Transfections were carried out using the FuGene6 transfection reagent (Promega, Madison, WI) according to the manufacturer's protocol. After 2 days, cells were harvested, and total RNA was extracted as previously described. The 3′ polyadenylation of RNA was performed using the Poly-A Polymerase Kit (Thermo Fisher Scientific, Waltham, MA). Reverse transcription was then carried out using the SuperScript III RT kit (Invitrogen, Waltham, MA) according to the manufacturer's instructions. miR-1247 siRNA:

mG/ZEN/mCmUmCmCmAmGmUmCmUmCmGmAmCmGmUmUmCmCmC/3ZEN/

1. Primer miRNA-1247-3p Forward: CGGGAACGTCGAGACTGGAGC
2. Primer miRNA-1247-5p Forward: ACCCGTCCCGTTCGTCCCCGGA
3. cDNA synthesis Adapter primer: GACGAGGACTCGAGCTCAAGCTTTTTTTTTTTTTTTTTT
4. Consensus Reverse Primer: GACGAGGACTCGAGCTCAAGC Negative control (human, mouse, rat) we bought from Integrated DNA Technology (Coralville, IA).

Control siRNA sequence:

mG/ZEN/mCmGmAmCmUmAmUmAmCmGmCmGmCmAmAmUmAmUmGmG/3ZEN/

We performed three independent experimental replicates (n = 3) with control and miR-1247 specific siRNA to ensure the reproducibility of miR-1247-5p and miR-1247-3p knockdown effect.

**fig. S1**

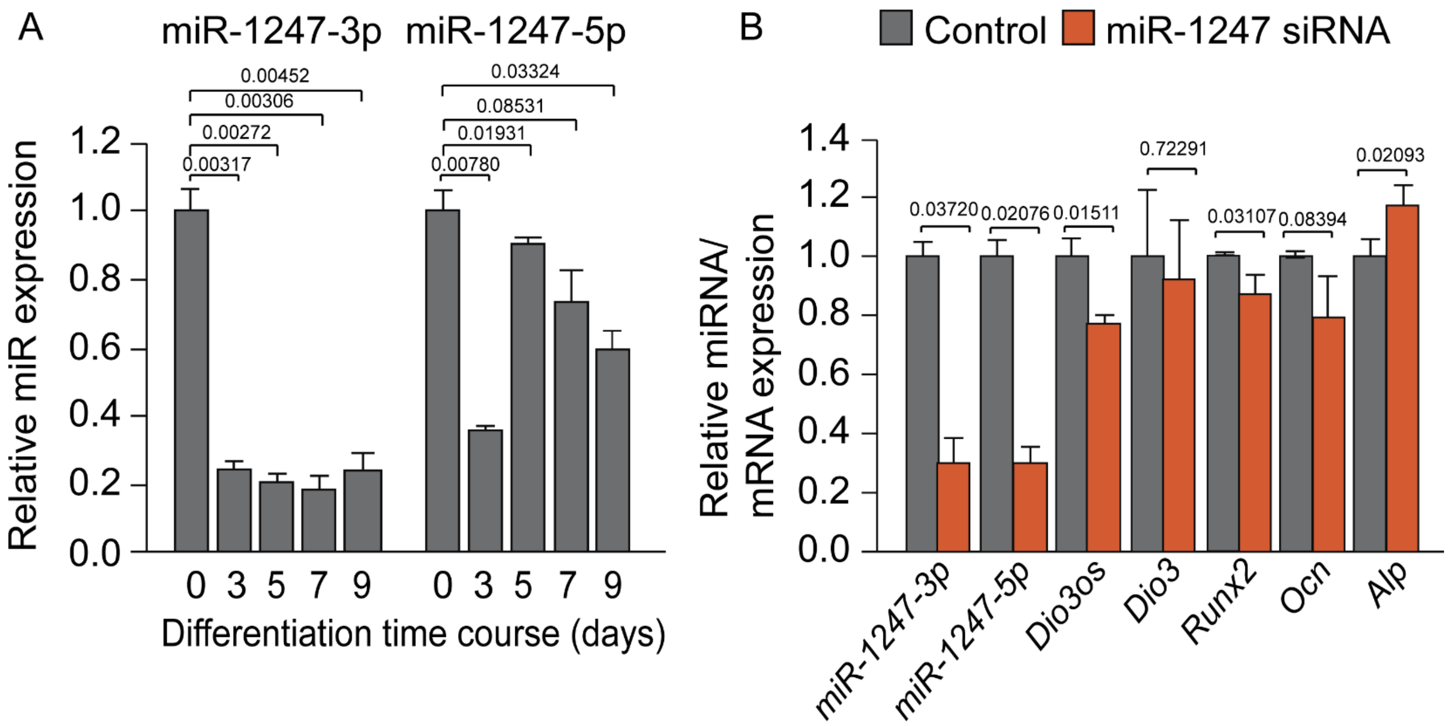

A
miR-1247-3p
miR-1247-5p
0.00452
0.00306
0.00272
0.00317
0.03324
0.08531
0.01931
0.00780
Relative miR expression
1.2
1.0
0.8
0.6
0.4
0.2
0.0
0 3 5 7 9
0 3 5 7 9
Differentiation time course (days)
B
Control
miR-1247 siRNA
Relative miRNA/ mRNA expression
1.4
1.2
1.0
0.8
0.6
0.4
0.2
0.0
0.03720
0.02076
0.01511
0.72291
0.03107
0.08394
0.02093
miR-1247-3p
miR-1247-5p
Dio3os
Dio3
Runx2
Ocn
Alp


**fig. S2**

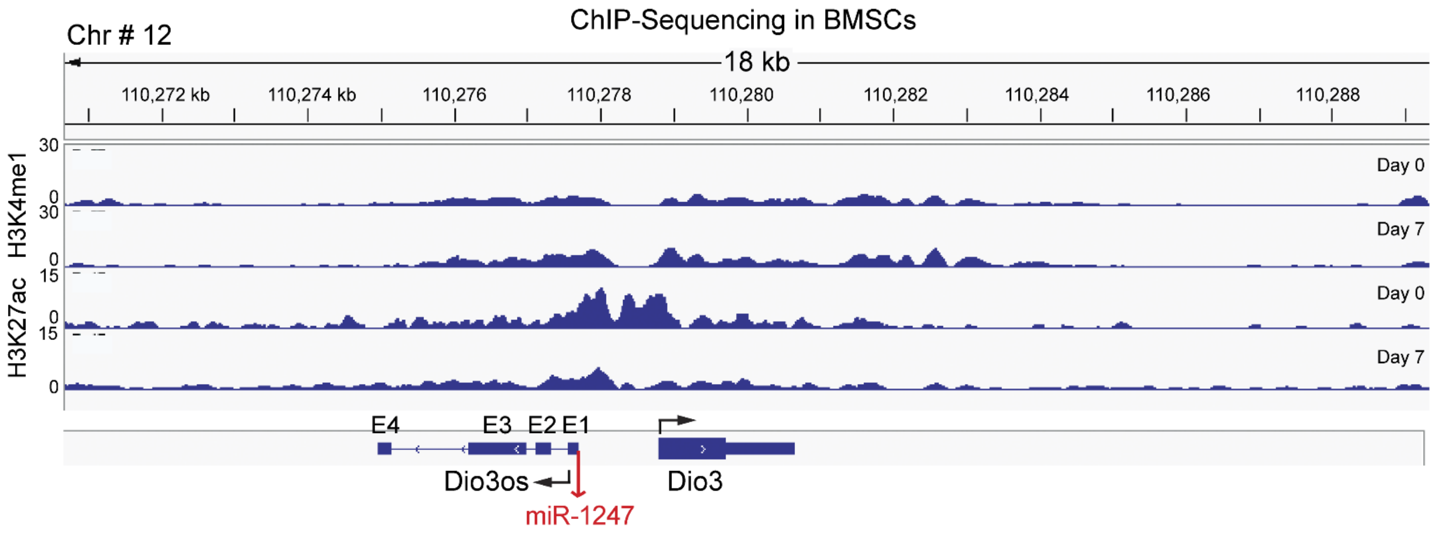

ChIP-Sequencing in BMSCs
Chr # 12
18 kb
110,272 kb
110,274 kb
110,276
110,278
110,280
110,282
110,284
110,286
110,288
H3K4me1
H3K27ac
30
0
15
Day 0
Day 7
E4
E3
E2
E1
Dio3os
Dio3
miR-1247

**fig. S3**

### A Schematic

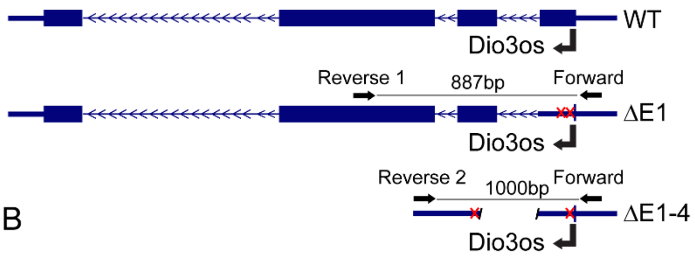


## B

| Primers | Sequence(5'-3') | PCR | PCR product | Genotype |
|---|---|---|---|---|
| Forward (F) | CTAGGGGTAGCTGTTGCCTG | F+R1 | 887 bp | WT |
| Reverse (R1) | GCCACTTTCACAGGCTGTTC | F+R1 | 700 bp | ΔE1 |
| Reverse (R2) | CCCCCAGGTAGGCTAGGATT | F+R1 | No product | ΔE1-4 |
| (F) and (R2) | | F+R2 | 1000 bp | ΔE1-4 |

## C

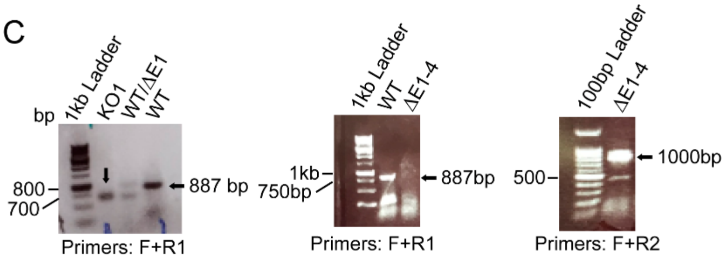


## D

Sequence ΔE1

5' TCTAAGCTCTCTCGCGCTTGCCTCGCCCGGCGCAGCCCCGGCCGCCGGGGGC
GGGCGCACCCGTCCCGTTCGTCCCCGGACGTTGCTCTCTGCCTTGGGTTCAGGG
GCGGCGGGGCTGCTTGACCTGGGGTCCTCGAGACCCTAAGCAAGTTCCACTTTA
GATACACCTCACAATGACCCCTCAGGTCAAGGCCTCAACCCGAAGGCAGCGCCTA
AGAATCGCACTCCCTAGAATGCTCCAGCCTCACAGGGGCCTTCTCTGGGGTAACG
GGACCCCAGCCGGATACTACAGCTCCAGCTGCTTGCGTCCCATGCATCGCCCTCT
CGGGAGAGACCAGCTACCTAACTTTCTCTACCAGAGACCGCGCTGGCGTCTCGGG
AGGGTGAGCACAGTCAGACCACTTTCCATTAGCCATGGAACTTGTCCGGAAAAGTC
GACCTCAAGTAGCCCAGAGGGCGCCCAGGTAGGCGCCCGCCCTCACCACCTGGT
TTCTGCTGTCCTGGCAGGATGCTCTGCCCAGGGAGTCCCTCGGTCGCGGGACCT
GTGGCCGCCAGAGCAGCTGGATCTGGAGTGTGCTGCCTGTGAACAGCCTGTGAA
AGTGGCA 3'

## E

Genomic Blast Results ΔE1

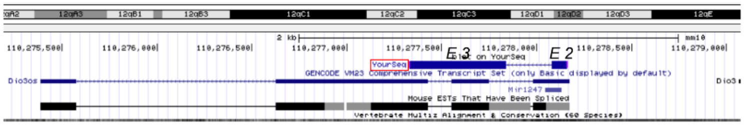


## F

Sequence ΔE1-4

5' TCTAAGCTCTCTCGCGCTTGCCTCGCCCGGCGCAGCCCCGGCCGCCGGGGGC
GGTAAGCCCGGAGGTACGAGGACGTGGCGCCCTGCAGCCTGACTGAGGTCATCT
AGGACTGGATCCGTGGAACATCCCCTTTTGGGTGTGGGATCTAACGAATATGCCCT
CCACCTGCCCACTTTATCCCGGTTCCTTTTCTAGGCTGTGCACCGCTCCCTCTGAA
ATAAGCGCTTATCCCTTTCCACTATTTTGAGAGTGCCCTTTGAAACACGTTTTCCCA
CCTCTGACTCAGGCCAACTAAGGTGGGCACAAAAATACTCTGAGGAGACAAGTCG
GCCAGTAGCTCAGTGATGATCTTGGGCTGATCTTGGAGAGCCAGGGTCTTAGCAT
CTTGACAATGTCTTCCTCATATGTTGCTCCACAGATGTAAAAGGCTCTAGACTCCTT
GCAACGAATTGCACTGCTCTTGTCTGCTACTCTGGGGTCCTGTAAAATCCTACCTG
TCCCCACCATCTGGCCATTTGCCTCCACTACTGTCTTACCTGACCTTTCCTGTCTTT
CCTTGAGGCCCCGCCCCCACCCTGCACCCTAACTTACGCTCTCACTGTGTCCACA
CATAACTAACCTCTCACTGTGGTCTCTGGTATTCTGCTAGCCCTGGGGGACAGGAC
AATGTGGGTACCTGGGGTCCTGACTGGTCTTGGGATGCAGGACTATGCATACAAGA
CAAATCCTAGACTGACCTGTGGGATGCAGGACTATGCATACAAGACAAATCCTAGCC
TACCTGGGGGGTTTA 3'

## G

Genomic Blast Results ΔE 1-4

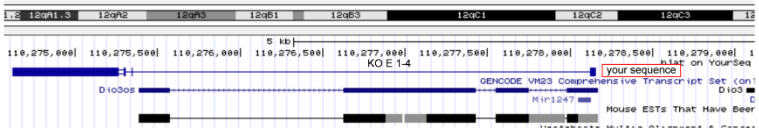

**fig. S4**

A

RNA-sequencing

| Sample Name | WT | ΔE1-4 |
|---|---|---|
| Total reads | 63783594 | 46361322 |
| Total maped reads | 61940710 | 44990108 |
| Uniquely mapped reads | 53829224 | 41283002 |
| Multiple mapped reads | 8111486 | 3707106 |
| Total mapping rate | 97.11% | 97.05% |
| Uniquely mapping rate | 84.39% | 89.05% |
| Multiple mapping rate | 12.72% | 8.00% |

B

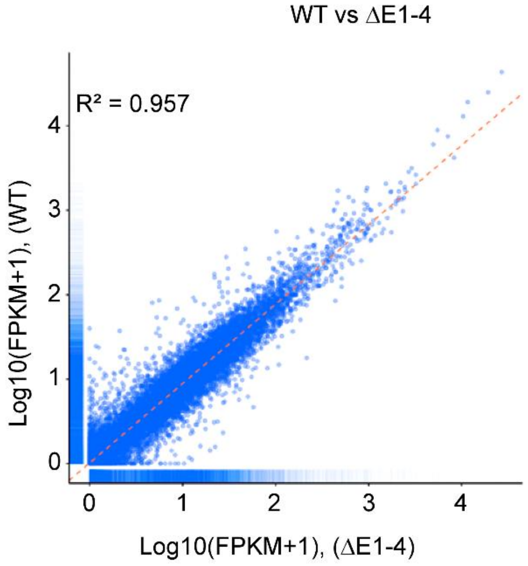


C

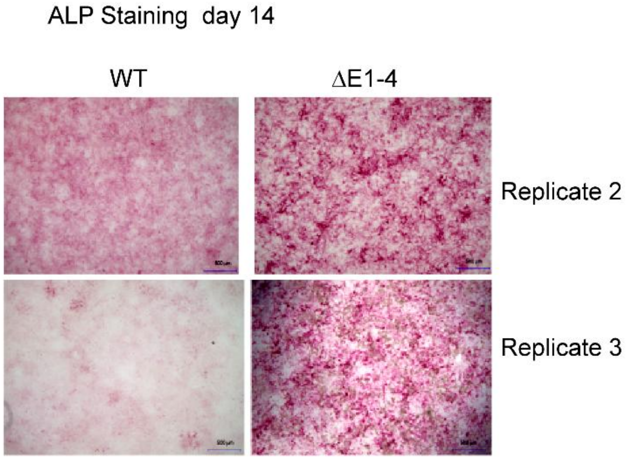


D

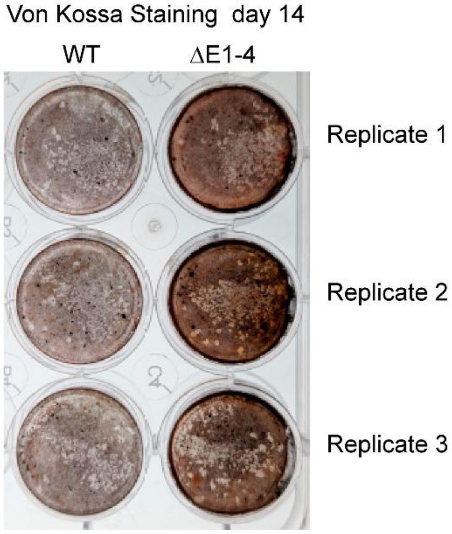

**fig. S5**

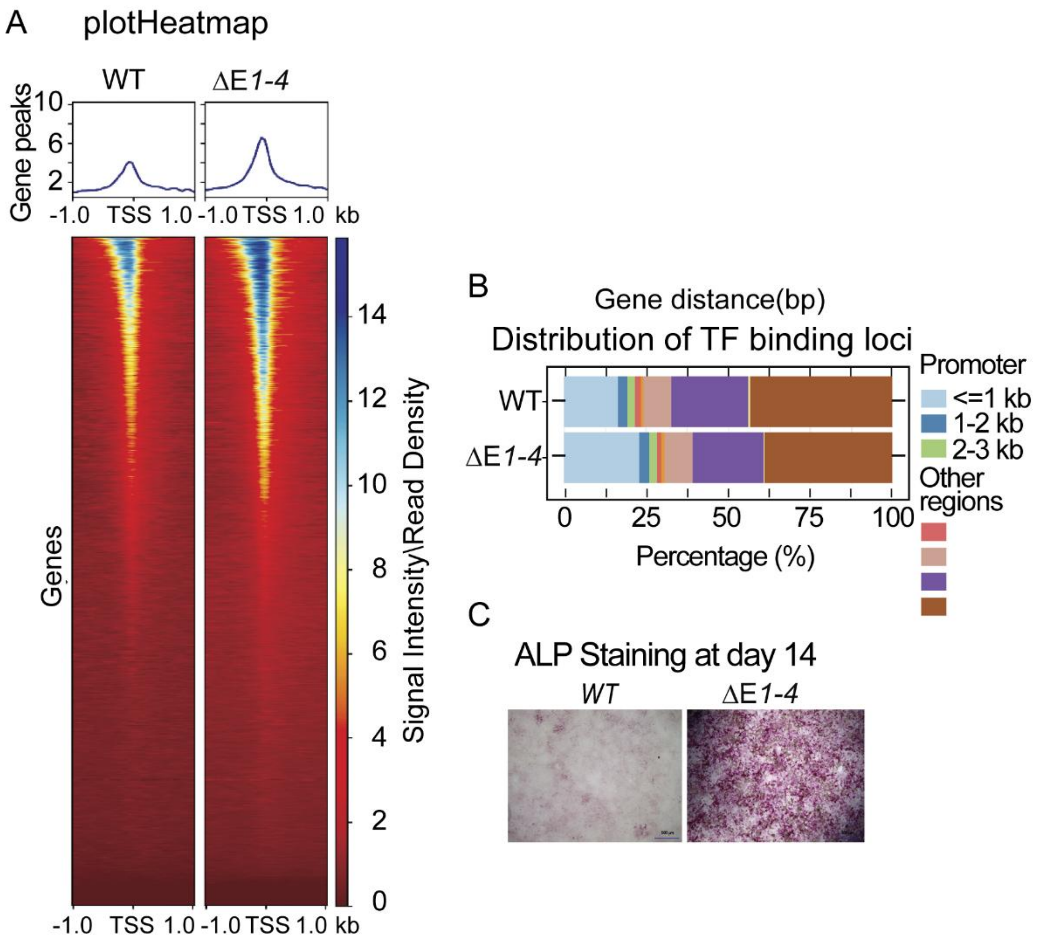
A
plotHeatmap
WT
ΔE1-4
Gene peaks
10
6
2
-1.0
TSS
1.0
kb
Genes
Signal Intensity\Read Density
14
12
10
8
6
4
2
0
B
Gene distance(bp)
Distribution of TF binding loci
WT
ΔE1-4
0
25
50
75
100
Percentage (%)
Promoter
<=1 kb
1-2 kb
2-3 kb
Other regions
C
ALP Staining at day 14
WT
ΔE1-4

fig. S6

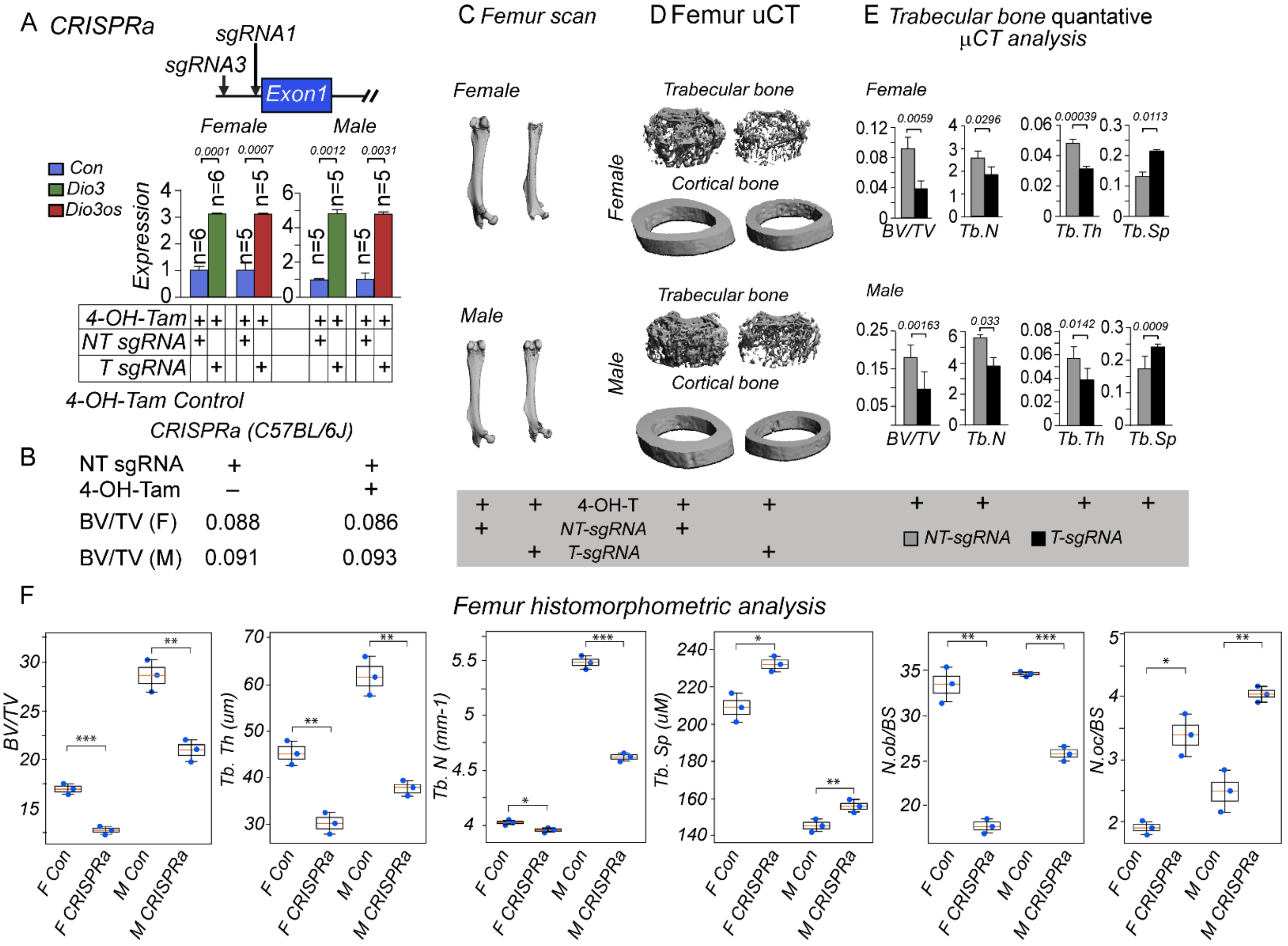
A CRISPRa
sgRNA1
sgRNA3
Exon1
Female
Male
Con
Dio3
Dio3os
Expression
4-OH-Tam
NT sgRNA
T sgRNA
4-OH-Tam Control
CRISPRa (C57BL/6J)
B
NT sgRNA + +
4-OH-Tam – +
BV/TV (F) 0.088 0.086
BV/TV (M) 0.091 0.093
C Femur scan
D Femur uCT
E Trabecular bone quantative µCT analysis
Trabecular bone
Cortical bone
BV/TV
Tb.N
Tb.Th
Tb.Sp
4-OH-T
NT-sgRNA
T-sgRNA
F
Femur histomorphometric analysis
BV/TV
Tb. Th (um)
Tb. N (mm-1)
Tb. Sp (uM)
N.ob/BS
N.oc/BS
F Con
F CRISPRa
M Con
M CRISPRa

**fig. S7**

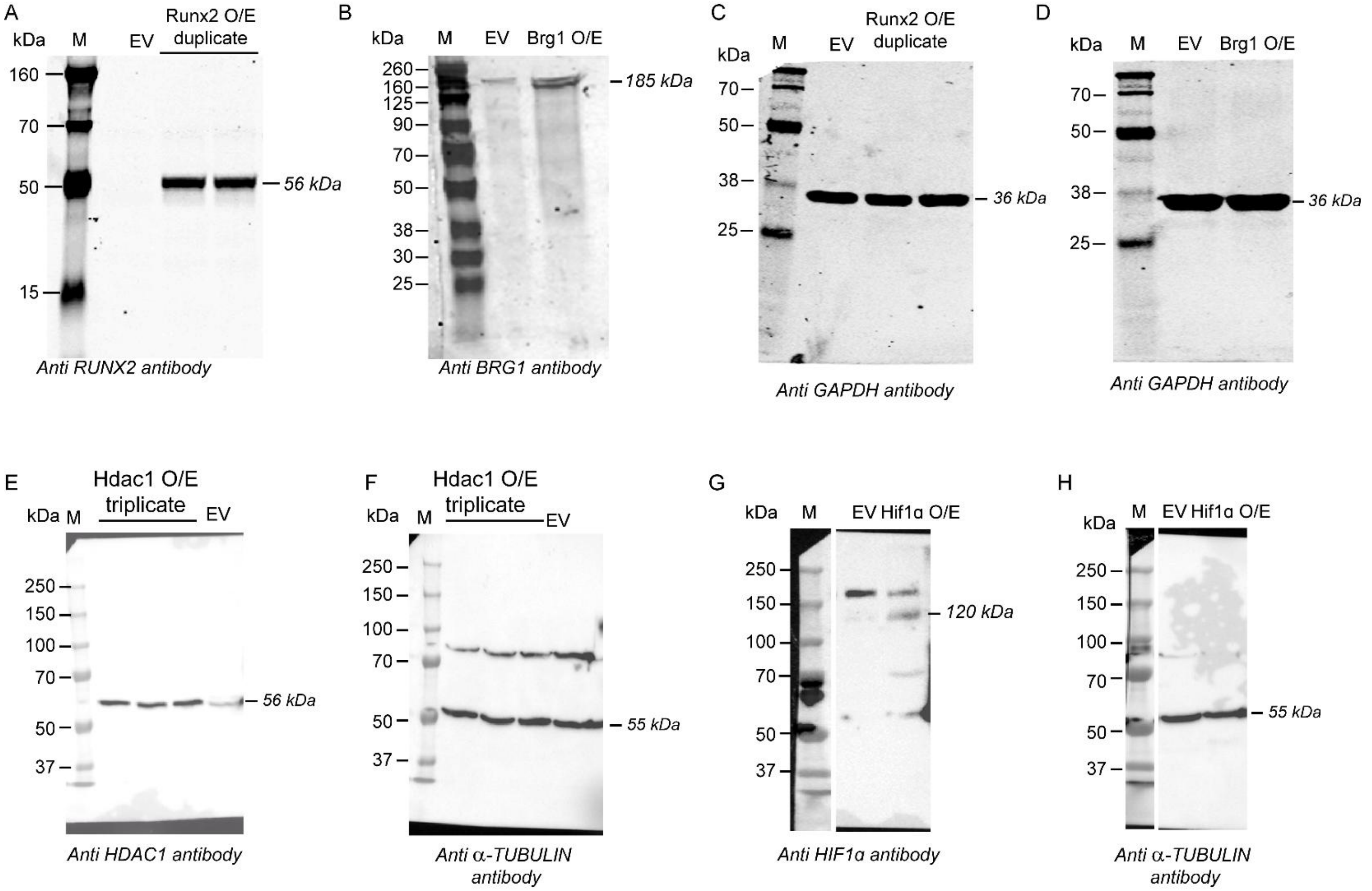


### Supplementary Figure Legends

**Figure S1**: **miR-1247 expression during MCM3T3-E1 differentiation and its effects on Dio3os, Dio3 and osteogenesis.** (A) MC3T3-E1 cells were differentiated in osteogenic media. Total RNA was isolated at different time points (0, 3, 5, 7, and 10 days). One microgram of total RNA was polyadenylated with PolyA Polymerase, reverse transcribed with oligo(dT) primer and amplified (RT-qPCR) by miR-1247-5p and 3p primers. The relative expression was normalized to U6 small RNA expression. (B) siRNA mediated knockdown of miR-1247 miRNA in MC3T3-E1 cells. MC3T3-E1 cells at 50-60% confluence were transfected with miR-1247 specific siRNA and non-specific control siRNA (100 nM). miRNA 1247-5p and 3p, Dio3, Dio3os, and osteogenic differentiation markers (Runx2, Ocn, and Alp) levels were assayed by RT-QPCR. Dark grey bars represent the control, and red bars indicate the siRNA knockdown. We performed three independent experimental replicates of osteoblast differentiation time points to ensure the reproducibility of our findings on miR-1247 expression. We also performed three independent replicates using both non-specific and specific siRNA to confirm the miR-1247 knockdown and its effect on Dio3os, Dio3, and bone marker genes.

**Figure. S2: Epigenetic profiling of enhancer-associated histone modifications across the Dio3os locus.** The analysis of ChIP-seq data for Bone Marrow Stromal Cells (BMSCs) using H3K4me1 and H3K27ac antibodies was performed across an 18 kb region of the Dio3os–Dio3 genomic area at days 0 and 7. The findings revealed that occupancies of the enhancer-associated marks (H3K4me1 and H3K27ac) were minimal within the intragenic exons 2–4 of Dio3os. In contrast, the promoter-proximal region exhibited strong enrichment of H3K4me3 and H3K27ac, indicating active promoter activity rather than that of an intragenic enhancer.

**Figure S3: Validation of CRISPR–Cas9–mediated Dio3os exon 1 and exon 1-4 deletions.** (A) Schematic representation of the Dio3os wild-type (WT) and knockout (KO) alleles showing the targeted deletion sites within exon 1 (KO E1) and exons 1–4 (KO E1–4). Arrows indicate primer positions (Forward, Reverse 1, and Reverse 2) used for genotyping. Red crosses display sgRNA-targeted genomic regions (B) Primer sequences and expected PCR product sizes for each genotype. (C) Representative agarose gel electrophoresis confirming successful generation of Dio3os knockout alleles. PCR with primers F + R1 yielded an ~887 bp product for WT, an ~700 bp product for KO E1, and no detectable band for KO E1–4. PCR with primers F + R2 generated a ~1000 bp product specific to KO E1–4 deletion. (D, F) Sanger sequencing of the KO E1 and KO E1–4 alleles showing the edited regions (remaining exon sequence highlighted in red). (E, G) Genomic BLAST alignment of the KO E1 and KO E1–4 sequences confirms precise localization of the deletions within the mouse Dio3os locus on chromosome 12 (positions 110,275,000–110,278,000 bp). Together, these data validate accurate CRISPR–Cas9 targeting and deletion of Dio3os exons used for subsequent functional studies.

**Figure S4**: **Verification of RNA-seq data quality and osteogenic differentiation following Dio3os knockout.** (A) Overview of RNA-seq mapping statistics for wild-type (WT) and Dio3os knockout (KO 1–4) osteoblasts. Both samples exhibited high sequencing quality with total mapping rates >97% and uniquely mapped read rates >84%, confirming robust library preparation and alignment efficiency. (B) The scatter plot comparing $\log_{10}$(FPKM + 1) values for all genes in WT versus KO 1–4 samples illustrates a strong correlation ($R^2 = 0.957$), indicating overall consistency of the transcriptome between samples and high reproducibility of the RNA-seq data. (C) Representative alkaline phosphatase (ALP) staining at day 14 in WT and KO 1–4 osteoblasts from two independent experiments demonstrating enhanced ALP activity and mineralization in Dio3os-deficient cells, consistent with increased osteogenic differentiation observed in RNA-seq analyses due to Dio3os CRISPR-Cas9-mediated deletion. Scale bars represent 100 μm.

**Figure S5: ATAC sequencing verified globally enhanced chromatin accessibility in bone-specific genes in Dio3os knockout cells** (A) Top panel showing peaks annotated to the nearest transcription start site (TSS) of genes or regulatory elements in the promoter when compared Dio3os knockout, over control MC3T3-E1 cells. Bottom panel: Heatmaps with transposase hypersensitive sites (THSs) of Dio3os regulated promoters in control, and Dio3os knockout MC3T3-E1 cells. The regions in the heatmaps are ranked from the highest ATAC-seq signal (top) to the lowest (bottom) and are aligned to the mm10 mouse genome. (B) Barplot showing the accessibility distribution of reproducible peaks of all seven regions. The bottom scale shows the percentage of open chromatin within 1-3 kb promoter and other regions of a gene. (C) Alp staining in control, and Dio3os knockout MC3T3-E1 cells at day 14 of osteoblast differentiation. ATAC-sequencing at day 14, we prepared two independent ATAC-Seq libraries (control and CRISPR knockout cells) along with their corresponding input libraries, made from DNA treated with Tn5 transposase. Y-axis scale (right) indicates signal intensity or read density of the transcriptional start sites.

**Figure S6: Phenotypic characterization of OB-specific CRISPR activation (CRISPRa) mice to study bone formation ( 2 month).** (A) Location of the CRISPRa sgRNAs. (A, lower panel) Dio3 and Dio3os mRNA quantification 4 weeks post-injection in femur bone lysates isolated from mice injected with 4-OH- Tamoxifen +/- non-targeting or targeting sgRNAs. CRISPRa: non-targeting sgRNA (n=6, n=5 female & male per group). Targeting sgRNA (n=6 female & n=5 male per group). (B) Femur bone BV/TV ratio from uCT values of CRISPRa female (F) and male (M) 2-month-old mice +/- 4-OH-Tamoxifen. (C) (upper and lower panels, female & male) Longitudinal microCT scans from CRISPRa + non-targeting sgRNA (NT-sgRNA) (left) and targeting sgRNA (T-sgRNA) (right) mice. Femoral trabecular and cortical microCT scans of one-month-old female (D upper panel) and male (D lower panel) from CRISPRa + NT-sgRNA (left) and T-sgRNA (right) mice. MicroCT quantitation of femoral trabecular bone of female (E upper panel) and male (E lower panel) from CRISPRa + NT-sgRNA (grey bar) and T-sgRNA (black bar) and mice. Representative graphs illustrate bone volume in relation to tissue volume, trabecular number, thickness, and spacing. Mice were sacrificed at 2 month of age after receiving 10 mg/kg/day of 4-OH-TAM (I/P) and sgRNA lentivirus (109 virus particles/kg/day, I/V) for four consecutive days in one month. Quantitation data here comply with CRISPRa, with n = 6 (T-sgRNA) and n = 5 (NT-sgRNA) female mice and n = 5 (T-sgRNA) and n = 5 (NT-sgRNA) male mice. P values: ns $P > 0.05$, * $P \leq 0.05$, ** $P \leq 0.01$, *** $P \leq 0.001$, **** $P \leq 0.0001$. 4-OH-Tam: 4-hydroxytamoxifen. NT-sgRNA: non-targeting sgRNA, and T-sgRNA: targeting sgRNA. (F) CRISPRa (gain-of-function) femur bone histomorphometry analysis showing reduced bone mass and enhanced bone resorption in female and male mice. Plastic sections of femurs from 8-week-old Control (non-targeting *sgRNA* + Tamoxifen, n=3 female & n=3 male per group) and CRISPRi (targeting sgRNA + Tamoxifen, n=3 female & n=3 male per group) mice were analyzed for BV (bone volume) /TV (total volume); Tb. N, trabecular number; Tb.Th, trabecular thickness; and Tb. Sp, trabecular space. Data are presented as individual data points with box plots and mean ± SEM (n = 3 per group). Statistical significance was determined using Welch's ttest comparing Control and experimental groups within each sex. Significance levels are indicated as follows: $p < 0.05$, ** $p < 0.01$, *** $p < 0.001$.

**Figure S7: Uncropped and raw immunoblot validation of transcription factor and chromatin regulator overexpression in MC3T3-E1 cells.** (A–H) Uncropped, raw Western blot images showing overexpression (O/E) of RUNX2 (A), BRG1 (B), HDAC1 (E), and HIF1α (G) in MC3T3-E1 osteoblast cells 24 h after transfection, compared with empty vector (EV) controls. Duplicate (RUNX2, BRG1, HIF1α) or triplicate (HDAC1) samples are shown as indicated. Corresponding loading controls are GAPDH (C, D) or β-tubulin (F, H). Total cellular lysates were prepared in 1X RIPA buffer containing protease inhibitors, resolved by 10% SDS–PAGE, transferred to PVDF membranes, and probed with antibodies specific for RUNX2, BRG1, HDAC1, HIF1α, GAPDH, or α-tubulin. Molecular weight markers (kDa) are indicated. These blots confirm overexpression of the indicated proteins under the conditions used for functional Luciferase promoter activity assays.
